# Uncovering the core microbiome and distributions of palmerolide in *Synoicum adareanum* across the Anvers Island archipelago, Antarctica

**DOI:** 10.1101/2020.02.20.958975

**Authors:** Alison Murray, Nicole Avalon, Lucas Bishop, Karen W. Davenport, Erwan Delage, Armand E.K. Dichosa, Damien Eveillard, Mary L. Higham, Sofia Kokkaliari, Chien-Chi Lo, Christian S. Riesenfeld, Ryan M. Young, Patrick S.G. Chain, Bill J. Baker

## Abstract

Polar marine ecosystems hold the potential for bioactive compound biodiscovery, based on their untapped macro- and microorganismal diversity. Characterization of polar benthic marine invertebrate-associated microbiomes is limited to few studies. This study was motivated by our interest in better understanding the microbiome structure and composition of the ascidian, *Synoicum adareanum*, in which the bioactive macrolide that has specific activity to melanoma, palmerolide A (PalA), was found. PalA bears structural resemblance to a combined nonribosomal peptide polyketide, that has similarities to microbially-produced macrolides. We conducted a spatial survey to assess both PalA levels and microbiome composition in *S. adareanum* in a region of the Antarctic Peninsula near Anvers Island (64° 46'S, 64° 03'W). PalA was ubiquitous and abundant across a collection of 21 ascidians (3 subsamples each) sampled from seven sites across the Anvers Island archipelago. The microbiome composition (V3-V4 16S rRNA gene sequence variants) of these 63 samples revealed a core suite of 21 bacteria, 20 of which were distinct from regional bacterioplankton. Co-occurrence analysis yielded several potentially interacting subsystems and, although the levels of PalA detected were not found to correlate with specific sequence variants, the core members appeared to occur in a preferred optimum and tolerance range of PalA levels. Taking these results together with an analysis of biosynthetic potential of related microbiome taxa indicates a core microbiome with substantial promise for natural product biosynthesis that likely interact with the host and with each other.

## 1. Introduction

Microbial partners of marine invertebrates play intrinsic roles in the marine environment at both the individual (host survival) and community (species distribution) levels. Host-microbe relationships are mediated through complex interactions that can include nutrient exchange, environmental adaptation, and production of defensive metabolites. These functional interactions are tied to the structural nature (diversity, biogeography, stability) of host and microbiome, and the ecological interactions between them. Studies of sponges, corals, and to a lesser degree, ascidians have revealed strong trends in invertebrate host species specificity to particular groups of bacteria and archaea. These studies have documented an underlying layer of diversity (e.g., [1–3]) in which habitat and biogeography appear to have strong influences on the microbiome structure and function [4–6].

The vast majority of host-microbiome studies have been conducted at low- and mid-latitudes from coastal to deep-sea sites. High latitude benthic marine invertebrate-associated microbiome studies are currently limited to the Antarctic, where just the tip of the iceberg has been investigated for different host-microbe associations [7] and ecological understanding is sparse. Antarctic marine invertebrates tend to have a high degree of endemicity at the species level, often display circumpolar distribution, and in many cases have closest relatives associated with deep-sea fauna. Whether endemicity dominates the microbiomes of high latitude benthic invertebrate is currently not known, nor is the extent of diversity understood within and between different host-associated microbiomes. Likewise, reports of core (conserved within a host species) microbiomes within Antarctic invertebrate species are sparse.

The few polar host-associated microbiome studies to date have documented varying trends in host-species specificity, with generally low numbers of individuals surveyed. For example, low species-specificity was reported in sponge microbiome compositions between different sub-Antarctic and South Shetland Island Mycale species [8] which shared 74% of the OTUs, possibly representing a cross-*Mycale* core microbiome. On the contrary, high levels of microbiome-host species specificity and shared core sequences within a species was observed in five McMurdo Sound sponge species [9]. The same was found across several Antarctic continental shelf sponge species [10]. Webster and Bourne [11] also found conserved bacterial taxa dominated by microorganisms in the class Gammaproteobacteria across the soft coral, *Alcyonium antarcticum*, sampled at three sites in McMurdo Sound. Another cnidarian, the novel ice shelf burrowing sea anemone *Edswardsiella andrillae*, contained novel microbiota, though the composition across a limited set of individuals was only moderately conserved in which some specimen were dominated by an OTU associated with the phylum Tenericutes and others, a novel OTU in the class Alphaproteobacteria [12]. Lastly, a single representative of the Antarctic ascidian *Synoicum adareanum* revealed limited rRNA gene sequence diversity, including the phyla Actinobacteria, Bacteroidetes, several Proteobacteria, Verrucomicrobia, and TM7 [13], though persistence of these taxa across individuals was not studied.

*Alcyonium antarcticum* (= *A. paessleri*) and *Synoicum adareanum* are both reported to be rich in secondary metabolites. The soft coral *A. antarcticum* produces sesquiterpenes that are unusual in bearing nitrate ester functional groups [14], while the ascidian *S. adareanum* is known to produce a family of macrolide polyketides, the palmerolides, which have potent activity against melanoma [15]. The role of the microbial community in contributing to host defensive chemistry, microbe-chemistry interactions and niche optimization, as well as microbe-microbe interactions, are unknown in these high latitude environments.

Here we have designed a study to investigate whether a core microbiome persists among *S. adareanum* holobionts that may inform our understanding of palmerolide origins. We conducted a spatial survey of *S. adareanum* in which we studied coordinated specimen-level quantitation of the major secondary metabolite, palmerolide A (PalA) along with the host-associated microbiome diversity and community structure across the Anvers Island archipelago (64° 46'S, 64° 03'W) on the Antarctic Peninsula (Figure 1). The results point to a core suite of microbes associated with PalA-containing *S. adareanum*, distinct from the bacterioplankton, which will lead to downstream testing of the hypothesis that the PalA producer is part of the core microbiome.

**Figure 1.**
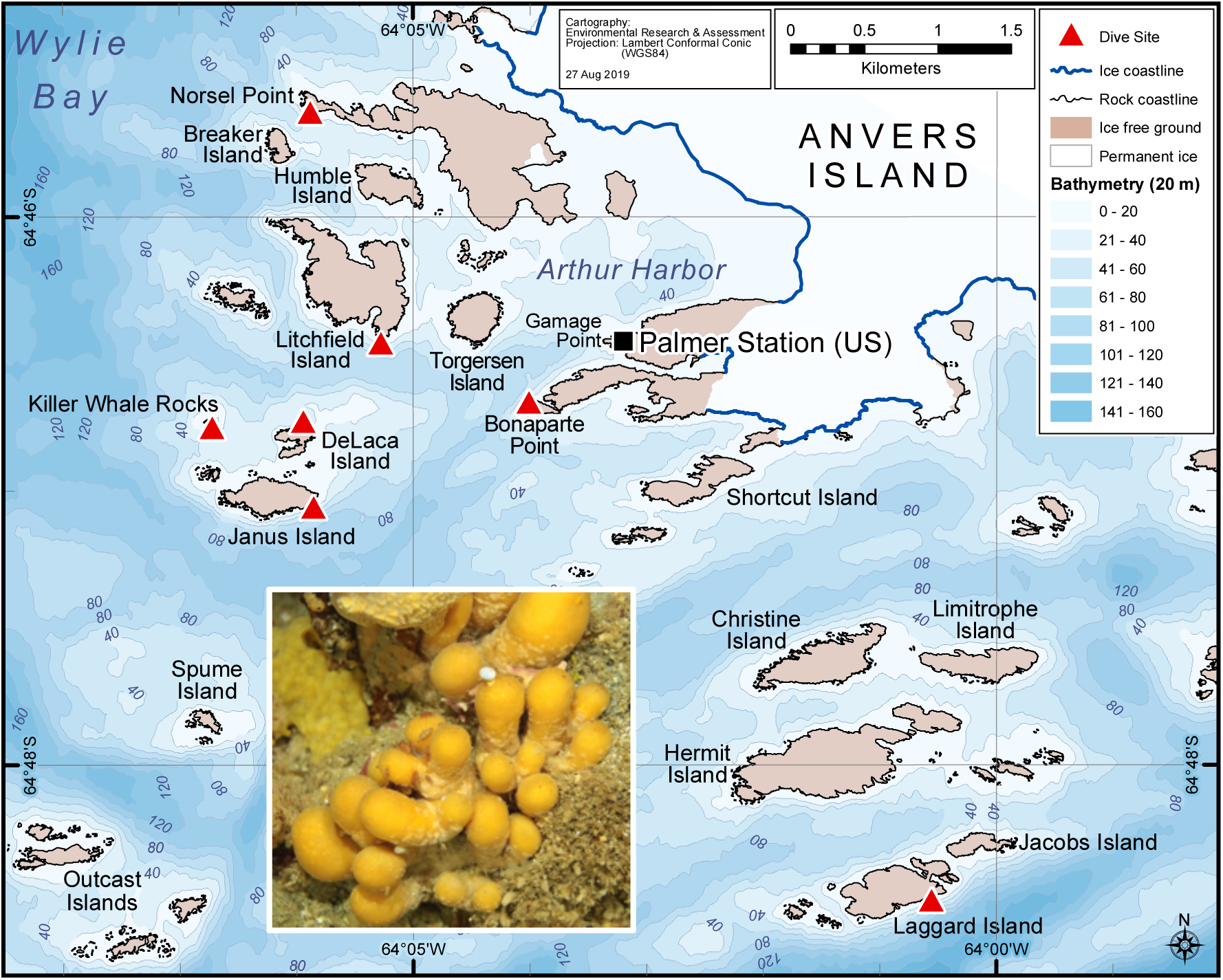
Bathymetric map of the study area off Anvers Island. *Synoicum adareanum* collection sites are shown with red triangles. The map was generated by Environmental Research & Assessment, Cambridge, UK, using Arthur Harbor bathymetry data from the PRIMO survey project 2004-06 (Dr. Scott Gallagher and Dr. Vernon Asper). Inset: Colonial ascidian, *S. adareanum*, which occurs in clusters of multiple lobes connected by a peduncle which together comprise a colony on the seafloor, collected at depths ranging from 24-31 m. Sampling site abbreviations in text: Bonaparte Point (Bon); DeLaca Island (Del); Janus Island (Jan); Killer Whale Rocks (Kil); Laggard Island (Lag); Litchfield (Lit); Norsel Point (Nor).

## 2. Results

### 2.1 Variation in holobiont PalA levels across ascidian colonies and collection sites

Typical procedures for natural products chemistry samples utilize bulk specimen collections for chemistry extraction (~ 30 individual ascidian lobes per extraction in the case of *S. adareanum*). Prior to this study, variation in PalA content at the individual lobe or colony level (inset, Figure 1) was unknown. Our sampling design addressed within and between colony variation at a given sampling site, as well as betwen site variation. The sites were constrained to the region that was logistically accessible by zodiac in the Anvers Island Archipelago off-shore of the United States Antarctic Program (USAP)-operated Palmer Station. *S. adareanum* colonies were sampled across seven dive sites (Figure 1) in which three lobes per multi-lobed colonies were sampled at three colonies per dive site, totaling 63 lobes for PalA comparison. PalA stands out as the dominant peak in all LC/MS analyses of the dichloromethane:methanol soluble metabolite fraction of all samples analyzed (e.g., Figure 2a). The range in PalA levels varied about an order of magnitude 0.49 −4.06 mg PalA per g host dry weight across the 63 lobes surveyed. Our study design revealed lobe-to-lobe, intra-site colony level and some site-to-site differences in PalA levels (p < 0.05) in the archipelago (Figures 2b, S2). Within a given colony, the lobe-to-lobe variation was often high and significantly different in 17/21 colonies surveyed. Significant differences in PalA levels between colonies were also observed within some sites (Janus Island (Jan), Bonaparte Point (Bon), Laggard Island (Lag), and Litchfield Island (Lit); see Figure 1) in which at least one colony had significantly different levels compared to another colony or both. Despite this, we found differences between some of the sites. Namely Bon was significantly lower than all six other sites. This site is the closest to the largest island, Anvers Island, and Palmer Station. Samples from Killer Whale Rocks (Kil) and Lit sites were also found to have significantly higher PalA levels than Jan, Bon and Norsel Point (Nor), although these did not appear to have a particular spatial pattern or association with sample collection depth.

**Figure 2.**
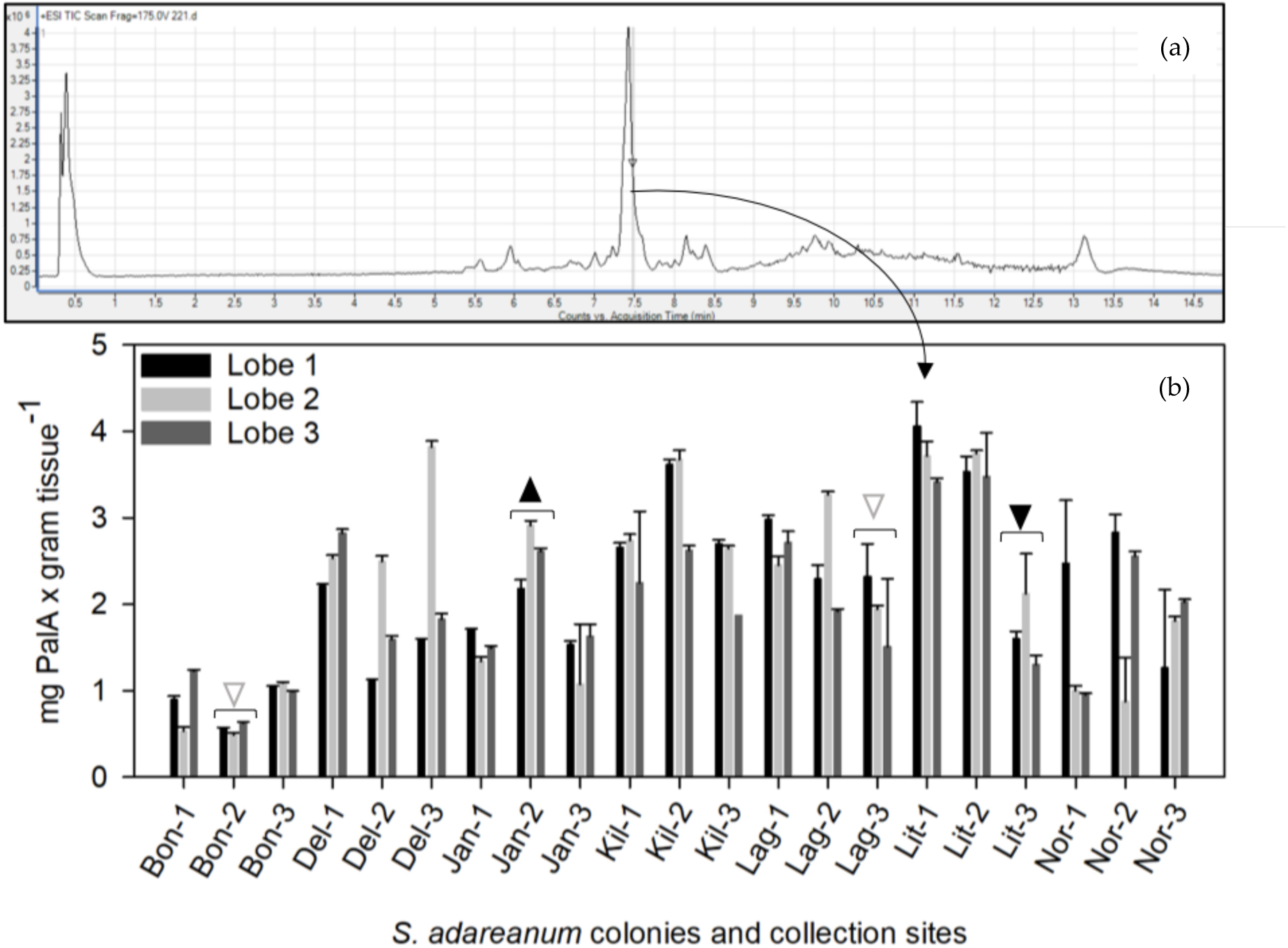
Palmerolide A (PalA) detection in *S. adareanum* holobionts. (a) Mass spectra derived from sample Lit-1a. The PalA peak dominates the dichloromethane-methanol fraction of the *S. adareanum* extract. (b) Levels of PalA normalized to tissue dry weight detected by mass spectrometry in *S. adareanum* holobiont tissues (three lobes per colony) surveyed in three colonies per site across the Anvers Island archipelago. Error represents individual lobe technical replication (standard deviation). Colonies with significant differences in PalA levels within a site are indicted with triangles, in which the direction of point indicates a significantly higher or lower colony. Filled triangles = significance (p < 0.05) in comparison to the other two colonies, and open triangles are those that were different from only one of the two colonies. Most colonies had significant lobe-lobe differences in PalA concentration, and some site-level differences were observed (Figure S2).

### 2.1. Characterization of host-associated cultivated bacteria

Given our interest in identifying a PalA producing microorganism, we executed a cultivation effort with *S. adareanum* homogenate on three different marine media formulations at 10 °C. 16S rRNA gene sequencing revealed seven unique isolates (of 16 brought into pure culture) at a level of > 99% sequence identity. All but one of the isolates was affiliated with the class Gammaproteobacteria, including five different genera commonly isolated from marine enviornments (*Shewanella*, *Moritella*, *Photobacterium*, *Psychromonas* and *Pseudoalteromonas*, of which nine were highly related). In many cases, we characterized their near neighbors as marine psychrophiles, many from polar habitats (Figure S1). The exception to this was the isolation of a cultivar associated with the Alphaproteobacteria class, *Pseudovibrio* sp. str. TunPSC04-5.I4, in which its two nearest neighbors were isolated from a marine sponge and a different ascidian. This result marks the first *Pseudovibrio* sp. cultivated from high latitudes. HPLC screening results of biomass from all sixteen isolates did not reveal the presence of PalA.

### 2.2 Synoicum adareanum microbiome (SaM)

To understand the nature of conservation in the composition of the host-associated microbiome of *S. adareanum* we identified the microbiome structure and diversity (based on the V3-V4 region of the 16S rRNA gene) with sections of the 63 samples used for holobiont PalA determinations. It resulted in 461 amplicon sequence variants (ASVs) distributed over 14 bacterial phyla (Table 1). The core suite of microbes, defined as those present in > 80% of samples (referred to as the Core80), included 21 ASVs (six of which were present across all 63 samples). The Core80 ASVs represented the majority of the sequenced iTags (95% on average across all 63 samples), in which the first four ASVs dominated the sequence set (Figure 3). The ASVs present in 50-79% of samples represented the Dynamic50 category and contained 14 ASVs, which represented only 3.3% of the data set. The remaining ASVs fell into the Variable fraction, which included 426 ASVs, representing 1.7 % of the iTag sequences, yet the majority of phylogenetic richness (Table 1). Comparative statistical analyses were conducted with the complete sample set, which was subsampled to the lowest number (9987) of iTags per sample. This procedure was limited by one sample (Bon1b) that underperformed in terms of iTag sequence yield. After the elimination of this sample from the analysis, the 62-sample set had 19,003 iTags per sample with a total of 493 ASVs, the same 21 sequences in the Core80 (with seven common to all 62 samples), the same 14 Dynamic50 sequences, and a total of 458 Variable ASVs.

**Table 1.** Relative proportions (average +/− standard deviations, n=63) of Phyla (and class for the Proteobacteria) across the different microbiome fractions.

| Phyla or Class | Whole Community | Core80 | Dynamic50 | Variable |
| --- | --- | --- | --- | --- |
| Proteobacteria |  |  |  |  |
| Gammaproteobacteria | 71.99 ± 6.64 | 73.28 ± 6.33 | 51.3 ± 23.16 | 43.71 ± 23.63 |
| Alphaproteobacteria | 22.9 ± 5.39 | 23.83 ± 5.93 | | 23.83 ± 19.7 |
| Deltaproteobacteria | 0.17 ± 0.1 | 0.16 ± 0.1 | | 1.11 ± 2.00 |
| Bacteroidetes | 2.83 ± 2.14 | 0.79 ± 0.69 | 46.55 ± 22.91 | 17.4 ± 14.69 |
| Verrucomicrobia | 1.56 ± 2.8 | 1.59 ± 2.93 | | 2.83 ± 3.97 |
| Nitrospirae | 0.27 ± 0.23 | 0.29 ± 0.24 | | 0.02 ± 0.17 |
| Planctomycetes | 0.12 ± 0.13 | | 2.15 ± 3.06 | 4.6 ± 12.84 |
| Actinobacteria | 0.1 ± 0.08 | 0.05 ± 0.05 | | 3.15 ± 3.53 |
| Patescibacteria | 0.02 ± 0.09 | | | 0.84 ± 2.05 |
| Dadabacteria | 0.02 ± 0.03 | | | 1.22 ± 2.03 |
| Uncl. Bacteria | 0.009 ± 0.017 | | | 0.72 ± 1.53 |
| Dependentiae | 0.004 ± 0.018 | | | 0.34 ± 1.99 |
| Chlamydiae | 0.002 ± 0.006 | | | 0.19 ± 0.83 |
| Acidobacteria | 0.000 ± 0.003 | | | 0.02 ± 0.13 |
| Chloroflexi | 0.000 ± 0.003 | | | 0.02 ± 0.13 |
| Epsilonbacteraeota | 0.000 ± 0.001 | | | 0.01 ± 0.07 |

**Figure 3.**
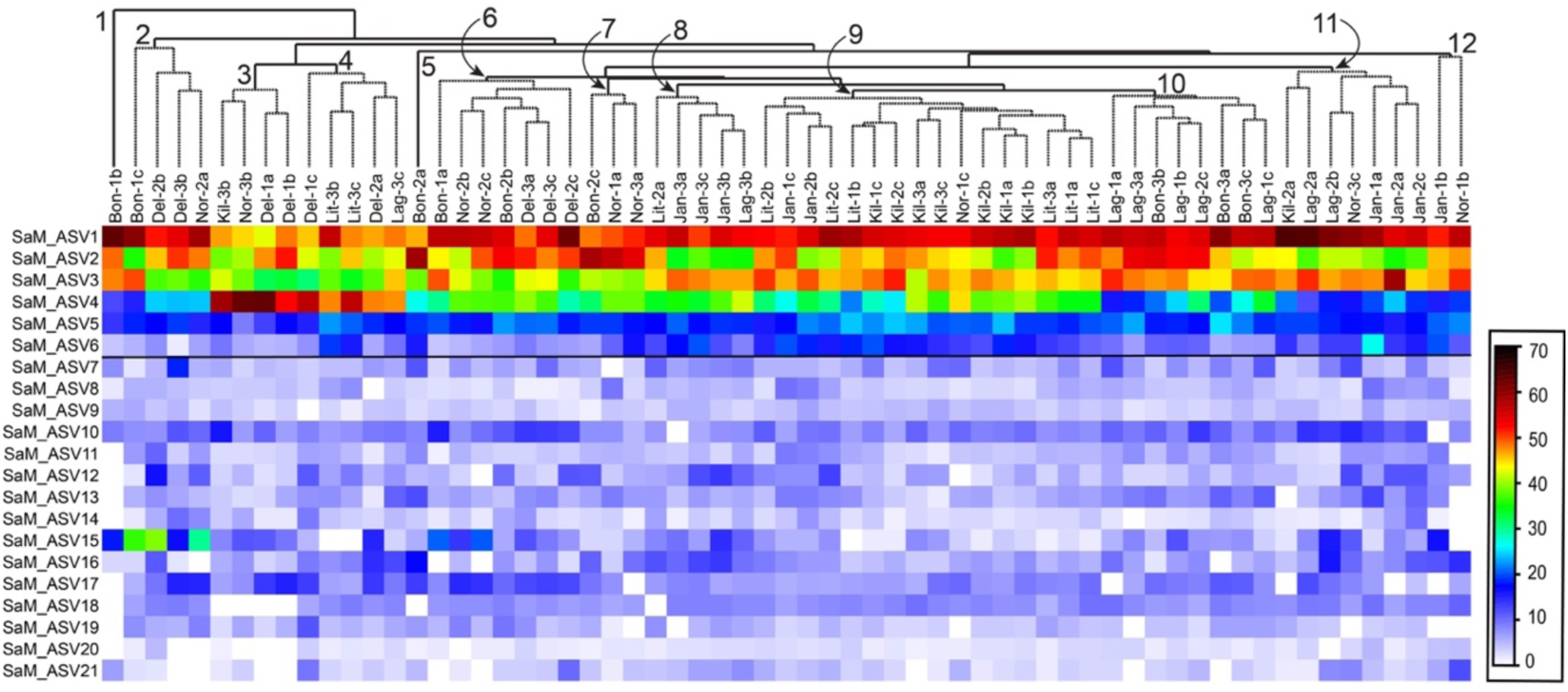
Heatmap of square root transformed ASV occurrence data for the core microbiome. ASVs (ranked and numbered) are shown on the y-axis, and 63-samples were hierarchically clustered, shown on the y-axis (site-based; square root transformed abundance data). Nodes with significant clusters are indicated from left to right (p < 0.05); order of clustering inside the node was not significant. The horizontal line drawn below SaM_ASV6 demarcates those ASVs that were present in all 63 samples.

ASV11 most closely matched a sequence in the Nitrospirae family from Arctic marine sediments. There were two Verrucomicrobium-affiliated sequences represented in different families (Puniceicoccaceae, SaM_ASV14, and Opitutacae, SaM_ASV15). Lastly, there were two ASVs affiliated with the phylum Bacteroidetes: one related to polar strain, *Brumimicrobium glaciale* (SaM_ASV19), and the other to a marine *Lutibacter* strain (SaM_ASV12).

Five Dynamic50 ASVs were affiliated with the marine Bacteroidetes phylum (Cryomorphaceae and Flavobacteriacae-related), in addition to six ASVs associated with the class Gammaproteobacteria (including four additional *Microbulbifer*-related sequences). There was also two additional phyla, an ASV related to a sponge-affiliated Verrucomicrobium isolate, and a Planktomycetes-related ASV (Table 1, Figure S1). A number of these ASVs were most closely related to isolates from marine sediments.

Interestingly, five sequences identified from earlier cloning and sequencing efforts with this host-associated microbiome (Figure S1; [13]) matched sequences in the Core80 and Dynamic50 data sets. Phylogenetic comparisons also revealed that the SaM isolates were distinct from the Core80 and Dynamic50 except for the *Pseudovibrio* sp. TunPSC04-5.I4 isolate, which was present in both the Core80, and the clone and sequencing study.

The hierarchical clustering of Core80 ASVs (based on Bray-Curtis similarity) across all 63 samples did not reveal strong trends in site or colony specific patterns (Figure 3). There were eight instances where 2 of 3 lobes paired as closest neighbors, and 4 of 8 primary clusters that included three lobes derived from the same colony. Sample Bon1b clustered apart from them all. In some cases, clusters could be attributed to specific ASVs. For example, cluster 2 (Figure 3) had the highest relative levels of SaM_ASV15 (an Opitutaceae family-affiliated sequence), whereas cluster 3 (Figure 3) had the highest relative levels of ASV4 (affiliated with *Microbulbifer* spp.).

Overall the community structures in the *S. adareanum* microbiomes across the 63 lobes surveyed had a high degree of similarity. Bray-Curtis pairwise similarity comparisons between lobes and colonies within each site were higher than 54% in all cases. When comparing the averages of pairwise similarity values within and between colonies, all sites, other than Lag, had higher similarity values within lobes in the same colony (ranging from 69.9 - 82.2%) compared to colonies within a site (66.5-81.0%; Figure S3), although the differences were small, and only Bon and Jan were significantly different (p< 0.05). We performed a two-dimensional tmMDS analysis based on Bray-Curtis similarity to investigate the structure of the microbiome between sites (Figure 4a). The microbiomes sampled at Kil and Lit had the highest overall degree of clustering (>75% similarity) while Kil, Lag, Lit, and Jan samples all clustered at a level of 65%. The microbiomes from DeLaca Island (Del) were the most dissimilar which was supported by SIMPER analysis in which two of the most abundant ASVs in the Core80 were lower than the average across other sites (SaM_ASV1 and 3) while others in the Core80 (SaM_ASV4, 15, and 17) were higher than the average across sites. We also performed a 2D tmMDS on the SaM fractions (Core80, Dynamic50, Variable) with and without permutational iterations which showed similar trends although partitioning of community structures between sites was more evident with the permutations (Figure S4). Site-based clustering patterns shifted to some degree in the different SaM fractions. For the Core80 alone, Jan samples clustered apart from the others. For Dynamic50, both Jan and Kil were outliers. Finally, for Variable fraction, Kil and Del samples clustered apart from the other sites. Variable fraction was more homogeneous, obscuring any site-to-site variability, while the core displayed tighter data clouds that showed a modest level of dispersion.

**Figure 4.**
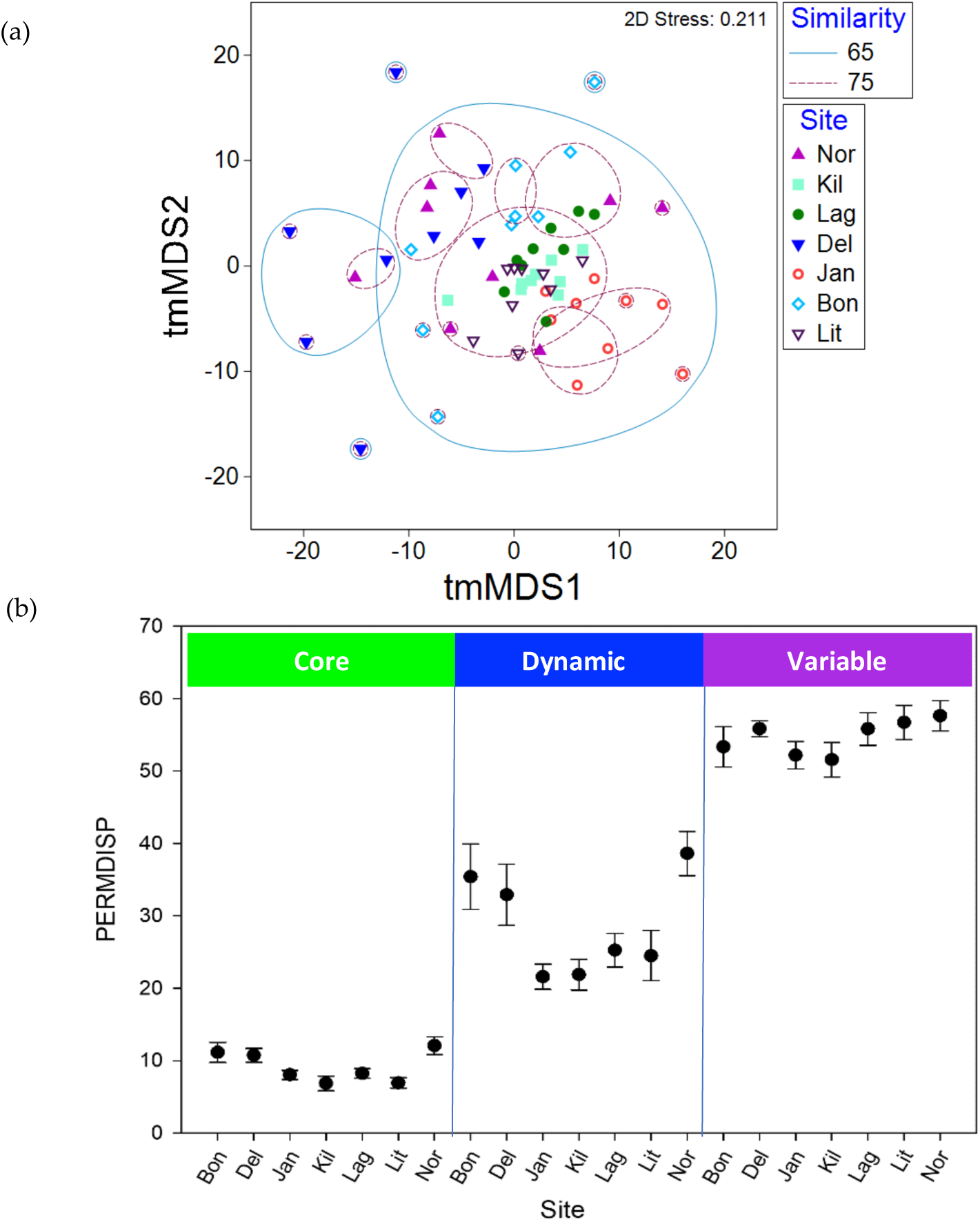
Similarity relationships amongst the *S. adareanum* microbiome samples in the Anvers Island archipelago. (a) tmMDS of Bray-Curtis similarities of square root transformed ASV occurrence data representing the microbiome of the 63 *S. adareanum* lobe samples using the complete ASV occurrence profiles. Microbiome samples with significant levels of similarity are shown (see legend). (b) B-diversity across Anvers Island archipelago sites represented by PERMDISP (9999 permutations) reveals differences between the highly persistent core, dynamic and variable portions of the *S. adareanum* microbiome (standard error shown). The degree of dispersion (variance) around the centroid changes significantly (p < 0.0001) for the different microbiome classifications, which the lowest levels of dispersion are found in the core microbiome.

A PERMANOVA analysis investigated the drivers of variability in community structure in which we tested the role of colony-level, site-level, and stochastic environmental variation. When evaluating the whole community (all sites and ASVs), site-to-site differences explained 25% of the variability in the microbiome (Table S1). Colony-to-colony differences explained 28%, while the remaining 47% of the variability was unexplained and is likely attributed to stochastic environmental variation. When the SaM fractions were analyzed separately, the most significant difference (p < 0.05) was in the Variable fraction of the microbiome in which the site-to-site differences explained only 19.2% of the variation and the residual (stochastic) level increased to 58.4% (Table S1). Conversely, PERMDISP (Figure 4), a measure of dispersion about the centroid calculated for each site (a measure of β-diversity), revealed differences in the community structures in each SaM fraction across sites, as well as differences between the microbiome fractions. The Core80 had a low level of dispersion (PERMDISP average of 9.1, range: 6.8-12.1), compared to the Dynamic50 (average 28.6, range: 24.5-38.6), which were moderately dispersed about the centroid, and were more variable with differences between the sites more apparent; Bon, Del, and Nor had higher values compared to the other four sites. The Variable fraction had high PERMDISP values (average 54.7, range: 51.6-57.6), representing high differences in β-diversity in which values were relatively close across the seven sites.

### 2.3 ASV co-occurrences, and relationship to PalA

To further investigate the ascidian-microbiome ecology, we performed a network analysis of the ascidian holobiont with a particular emphasis on ASV co-occurrence and ASV ecological niche associated with PalA. The co-occurrence network depicted a sparse graph with 102 nodes and 64 edges (average degree 1.255; diameter 15; average path length 5.81) that associated ASVs from similar SaM fractions (assortativity coefficient of 0.41), confirming the robustness of the above discrimination. Upon inspection of the co-occurrence network, a dominant connected component contained 42.1% of ASVs, in which we identified three highly connected modules (via the application of an independent network analysis WGCNA, soft threshold = 9, referred to here as subsystems; Figure 5). The three subsystems were not significantly associated with PalA (resp. Corr= −0.23; 0.033; −0.17 for subsystems 1, 2 and 3). The network also contained several smaller systems (2-6 nodes), and 30 singletons, only one of which was in the Core80 (Gammaproteobacteria class, *Endozoicomonas*-affiliated). The three subsystems included ASVs from the Core80, Dynamic50 and Variable SaM fractions, but were unevenly distributed. Mostly driven by Core80 SaM fractions, Subsystem 1 harbored most of the highly represented ASVs, including the *Microbulbifer*-related ASVs as well as the *Pseudovibrio* ASV. Subsystem 2, interconnecting Subsystems 1 and 3, is mostly driven by Variable fraction ASVs and includes lower relative abundant Core80 *Hoeflea* and the Opitutaceae ASVs, as well as several Bacteroidetes taxa. Finally, Subsystem 3 was smaller. It included several diverse taxa dominated by Dynamic50 fraction, but still including two representatives from the Core80, *Lentimonas*, and Cryomorphaceae-affiliated ASVs. The overall role of Variable ASVs in the network is surprising, as they appear to be critical nodes linking the Subsystems, via 13 edges that lie between Subsystems 2 and 3. Another, perhaps noteworthy smaller connected component is in the lower left of the graph in which three Core80 ASVs, including the chemoautotrophic ammonia and nitrite oxidizers, *Nitrosomonas* and *Nitrospira*, were linked along with an uncharacterized Rhodospiralles-related ASV, and a Variable ASV.

**Figure 5.**
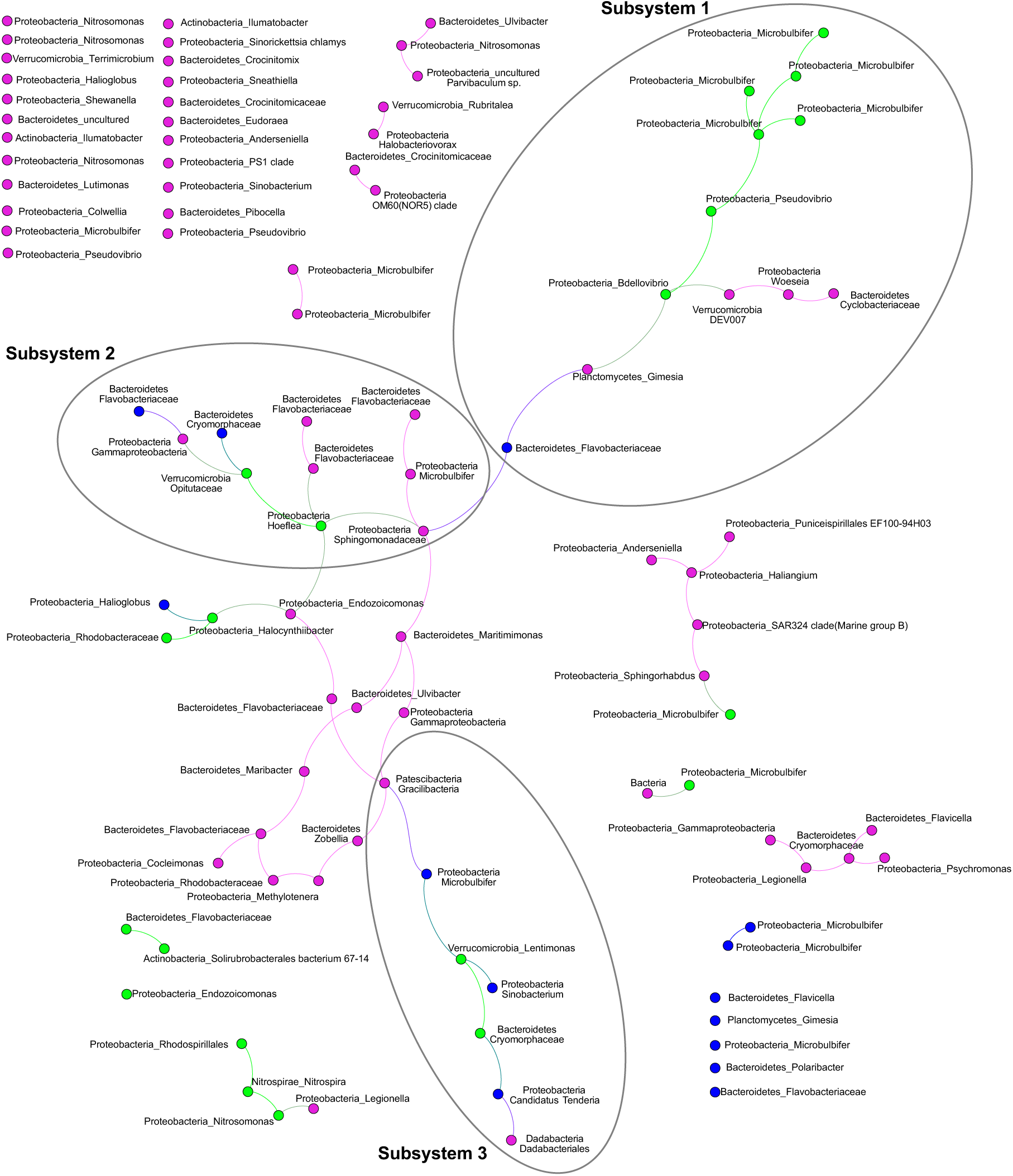
ASV co-occurrence network. The largest connected component of the co-occurrence network (seeded with ASVs found in at least 5 samples, 102 in total) identified three subsystems. Node colors represent the microbiome fractions (Core80, green; Dynamic50, blue; Variable, pink). Taxonomic identities of the ASVs are shown next to the nodes, with the phylum_highest level taxon identified shown.

To address the first-order question as to whether there was a relationship between the PalA concentration levels detected in the LC/MS analysis and the semi-quantitative ASV occurrences of *S. adareanum* ASVs, we performed three complementary analyses: correlation analysis, weighted co-occurrence networks analysis (WGCNA) and niche robust optima with PalA concentration as a variable. Pearson correlations between ASV occurrences and PalA concentrations ranged from −0.33 to 0.33 at the highest (24 of which were significant, ≤ 0.05; although none of those with a significant relationship were part of the core microbiome and were present in < 50% of the samples with a high occurrence of 24 sequences), suggesting little relationship at the gross level of dry weight normalized PalA and ASV relative abundance. This indicated that relative abundance of ASV is not a good predictor of PalA. A complementary WGCNA result showed no significant association between subsystem co-occurrence topology and PalA, pointing to the lack of relationship between microbial community structure and PalA. Finally, another aspect of the relationship between ASV occurrence and PalA levels was explored using the robust optimum method, which estimates the ecological optimum and tolerance range for ASVs about PalA. In this case we calculated the PalA niche optimum for each ASV and ranked them based on the median (Figure S5). Core80 ASVs showed a consistent PalA niche range. Furthermore, the median optima values of the Dynamic50 ASVs lie, for the most part, in lower or higher PalA optima compared to the Core80, with a substantial niche overlap between Core80. The Variable ASVs collectively lie at the lower and higher extremes of the optimum and tolerance range. Altogether, these complementary analyses advocated for not considering an individual nor a community effect on PalA, but rather, acclimation to the high levels of PalA observed that likely rely on unknown metabolic or environmental controls.

### 2.4 Culture collection:microbiome and bacterioplankton comparisons

To address whether the composition of the *Synoicum adareanum*-associated bacterial cultivars were also present in the SaM or in the free-living bacterioplankton (< 2.5-µm fraction), we considered the overlap in membership between the isolate 16S rRNA gene sequences and these other two data sets. Representation of isolates in the 16S rRNA gene ASV data set was estimated by comparing the two sequence data sets, albeit different ascidian samples were used for the culture collection and the *S. adareanum* survey. Excepting *Pseudovibrio* sp. TunPSC04-5I4 that was present in the Core80 with a 100% sequence match, two other isolates also had 100% matches to sequences in the Variable SaM (BONS.1.10.24 and BOMB.9.10.19). Three other isolate 16S rRNA sequences (BOMB.3.2.14, BOMB.9.10.16, BOMB.9.10.21) were found to match sequences in the Variable microbiome at a level of 97% or higher. Only the *Pseudoalteromonas*-related isolate and the *Shewanella*-related isolate sequences were not detected with relatives at a level of at least 97% identity to the SaM ASVs; which could be explained by under-sampling. The bacterioplankton composition was dominated by Gammaproteobacteria ASVs (47.35% of all ASVs; Table S2) in which six of the isolates (BOMB.9.10.21, BONS.1.10.24, BOMB.9.10.16, BOMB.3.2.20, BOMB.3.2.14, BONSW.4.10.32) matched sequences in the bacterioplankton data set at > 99.2% identity; even at a level of 95% sequence identity the remaining ten isolates did not match sequences in the plankton, including the *Pseudovibrio* sp. str. TunPSC04-5I4 isolate.

### 2.4 Microbiome:bacterioplankton comparisons

Although at a high taxonomic level Proteobacteria and Bacteroidetes phyla dominated the microbiome and bacterioplankton (Table S2), the relative proportions varied and the taxa represented were quite different. Membership between SaM and the bacterioplankton data set indicated a low-level overlap at 100% identity (Figure S6), with 39 of 604 perfectly matched ASVs. At 100% ASV sequence identity, the results indicate a single, Core80 SaM ASV was a perfect match with the bacterioplankton data set – the *Microbulbifer*-associated sequence that is the most abundant across all 63 SaM data sets. Interestingly, this sequence was only identified in one bacterioplankton sample (IPY-225-9) at a low occurrence (14 of > 1.18 million tags distributed across 604 ASVs). There were three (of 14) Dynamic50 ASVs that were perfect matches with the bacterioplankton ASVs. These were affiliated with a poorly classified Bacteroidetes family Flavobacteraceae ASV (77% of SaM samples), a Gammaproteobacteria-associated *Sinobacterium* (73% of SaM samples), and a second Gammaproteobacteria-associated ASV that is associated with *Candidatus* Tenderia (71% of SaM samples). The remaining 35 perfect match ASVs between the two data sets were classified as part of the Variable microbiome. These sequences fell across four phyla and nine classes, 13 of which were well-distributed across the Bacterioplankton data set samples (>50%).

At a level of 97% ASV sequence identity, there were three additional matches between the Bacterioplankton data set and the Core80. These included an Alphaproteobacteria-related *Hoeflea* and *Halocynthibacter*-related sequences, and a *Nitrospira*-related sequence. There were also two other Dynamic50-related ASVs: these were both related to unclassified Flavobacteriaceae. The rest (79 ASVs) of the matches at >97% were affiliated with the Variable fraction of the microbiome.

## 3. Discussion

This study reports our growing understanding of the microbiome composition of PalA-containing *S. adareanum*. To enhance our understanding of the ecology of the PalA-containing ascidian, *S. adareanum*, we investigated the ascidian colony microbiome and PalA chemistry levels at an individual lobe level, and compared the ascidian microbiome to the plankton. This comparison allowed us to address questions regarding variability in PalA content and whether a conserved core microbiome occurs across these PalA-containing Antarctic ascidians, thereby supporting the logic that if a microbial producer synthesizes PalA, the producing organism should be present in all PalA-containing *S. adareanum* samples. Before this study, however, we did not have quantitative data at the level of the individual ascidian lobe that forms the pedunculated *S. adareanum* colonies (Figure 1). This discussion focuses on the core microbiome, then takes a broader look at secondary metabolite distributions about the microbiome and in other marine invertebrates as well as the biosynthetic potential of core membership, and concludes with information gained from our initial cultivation effort.

### 3.1 Core microbiome

Ascidian (host)-microbiome specificity is an active area of research. Compared to sponges and corals, for example, ascidian microbiomes are less-well characterized. To get a broader perspective on the microbiome we also used cultivation-independent approaches. We found that the Antarctic ascidian *S. adareanum* has a persistent core microbiome across the Anvers Island archipelago that is distinct from the plankton. This dissimilarity between ascidian host-associated microorganisms and bacterioplankton appears to be a consistent observation across the global ocean (e.g., [16–19]). The Core80 is comprised of ASVs that numerically dominate the community, as well as those representing only a fraction of a percent of the sequences surveyed. Although ascidian symbioses have not yet been systematically studied in the Antarctic, better-studied lower latitude ascidian microbiomes provide several examples for comparison. The overall trend across ascidian microbiome studies to date suggests that there is a high degree of both geographical as well as host-species level specificity of microbiome composition (e.g., [16, 17, 20]). The same appears to be true of *S. adareanum.* Though this study was restricted to a small geographical region, we identified a conserved core of 21 16S rRNA gene sequence types across 63 individual pedunculate lobes studied. We attribute the detection of this high degree of persistent members in part to the uniform homogenization, extraction, and sequencing methodological pipeline applied. Microbiome analysis is sensitive to sequencing depth, quality parameter choices, and algorithmic differences in data-processing pipelines (amplicon sequence variants vs. cluster-derived operational taxonomic units), which can impact direct comparisons between studies. Along these lines, the numerous highly related Microbulbifer ASVs would have fallen into a single OTU (97% sequence identity), resulting in a core with 14 members. These limitations aside, our findings are in line with several other ascidian microbiome studies from lower latitudes in terms of the relative size of core membership (where core definitions vary to some degree between studies). For example, *Styela plicata*, a solitary ascidian, was reported to have a core membership of 10 [21] to 16 OTUs [22]. Other solitary ascidians, including Herdmania momus had a core of 17 OTUs [21], while two *Ciona* species ranged from 8-9 OTUs [23]. Temperate colonial ascidians *Botryliodes leachi* and *Botryllus schlosseri* ranged from 10-11 members in their core microbiomes [20]. Also, an extensive survey of 10 different ascidian microbiomes (representing both solitary and colonial forms) conducted on the Great Barrier Reef reported core memberships ranging from 2 to 35 OTUs [16], while the numbers of individuals surveyed in each case were only 2-3. Note that a few other studies reported much higher numbers of shared OTUs ranging from 93-238 [18,19]; the scale of sequencing was higher in these later studies. Further, as others have reported [24], the membership of these core ascidian microbiomes is distinct, and in the case of SaM, the core microbiome diversity appears to be unique at the ASV level, although several taxa are in common with other ascidian-associated microbes at the genus level including *Microbulbifer* associated with *Cystodytes sp.* [25], *Pseudovibrio* with *Polycitor proliferus* [26] and an *Endozoicomonas* specific-clade was identified in a survey of a number of ascidians [27].

Predicted metabolic abilities of the Core80 taxa suggest aerobic heterotrophy (aerobic respiration – organic carbon is the carbon and energy source), microaerophily (growth in low oxygen conditions) and chemoautotrophy (CO_2_ fixation provides carbon and reduced chemicals provide energy, e.g., NH_4_+ and NO_2_-) are themes amongst the Core80, in which the most abundant ASVs are high molecular weight carbon degraders. The *Microbulbifer* genus has members known to degrade cellulose [28], and perhaps non-coincidently, ascidians are the only known invertebrate capable of cellulose biosynthesis in the marine environment (e.g., [29, 30]). From this, we could speculate that the *Microbulbifer* strains associated with *S. adareanum* could occupy a commensal, if not somewhat antagonistic relationship [31]. In support of this possibility is the fact that the only overlapping sequence between the Core80 and the bacterioplankton was a *Microbulbifer* sequence, which was a rare sequence in the plankton – suggesting it may be an opportunistic member of the *S. adareanum* microbiome. Besides, free-living and sponge-associated isolates from the *Microbulbifer* genus have been found to produce bioactive compounds including pelagiomicins [32] and parabens [33], respectively. This observation, in the least, suggests that the *Microbulbifer*(s) is/are likely well-adapted to their ascidian host and might be considered a potential PalA producing organism.

The NH_4_+-oxidizing *Nitrosopumulis*-related Thaumarchaeota have been commonly detected in ascidian microbiomes [16, 21, 22, 34], which contrasts phylogenetically, but not in terms of overall function with the NH_4_+-oxidizing *Nitrosomonas*-ASV that was part of the SaM core. The niche however has been reported to be different for the archaeal and bacterial ammonia oxidizers in which the archaea tend to be found in oligotrophic systems, while the bacteria (e.g., *Nitrosomonas*) can tolerate high levels of dissolved ammonia [reviewed by 35]. This result might reflect both the environment, and in situ *S. adareanum* tissue ammonia levels where it may accumulate, as several studies have reported on the high levels of oxidizing ammonia Thaumarchaeota in the coastal waters of the Anvers Island archipelago. This group, however, is numerous only in winter to early spring waters [36–38], while this study was conducted with samples collected in Fall when the ammonia-oxidizing Thaumarchaeota are not abundant in the coastal seawater [36], advocating for the comparisons between the SaM with bacterioplankton collected in both summer and winter periods.

In a similar vein, although we did not intentionally conduct a temporal study, the data from samples collected in 2007 and 2011, appear to suggest that a number of the core microorganisms are stable over time. We found several ASVs in 2011 samples that matched (at 100% sequence identity) cloned sequences from samples collected in 2007. Stability of the ascidian microbiome over time has been reported in a few studies [17, 24, 34]. Studying the persistence of the core membership over the annual cycle would be interesting (and provide compelling evidence for stable relationships) in this high latitude environment where light, carbon production, and sea ice cover are highly variable.

The co-occurrence analysis indicated three subsystems of ASVs that co-occur within *S. adareanum*. A small side-network included the two taxa involved with the 2-step nitrification process, including the Nitrosomonas ASV mentioned above and a Nitrospira ASV. Even though at present, the functional underpinnings of the host-microbial system have not been studied, the co-occurrence relationships provide fodder for hypothesis testing in the future. One interaction network that warrants mentioning here is the ASVs in Subsystem 1, which harbor several Core80 *Microbulbifer* ASVs, and the *Pseudovibrio* ASV are linked to a *Bdellovibrio* ASV that is also a member of the Core80. Members of the Bdellovibrio genus are obligate bacterial predators [32] that penetrate the outer membrane and cell wall of their prey. The linkage position in the subsystem is compelling in the sense that the *Bdellovibrio* could potentially control the abundance of the “downstream” less connected members of the network. Lastly, the positions of a couple of Dynamic and several the Variable ASVs in the network, as links between the subsystems, was unexpected. The central positions of these ASVs suggest that they may not be merely stochastic members of the microbiome; that they could play opportunistic, adaptive or ecological roles in the functionality of the microbiome subsystem(s) which potentially participate in different aspects of the holobiont system in particular, by promoting the switch between different ecological modes supported by different subsystems. Such roles were proposed for dynamic members of the *Styela plicata* microbiome [21].

The culturing effort succeeded in isolating a *Pseudovibrio* strain that is a crucial member of the Core80. In addition, several other Gammaproteobacteria-affiliated strains which matched sequences in the Variable SaM and the bacterioplankton were cultivated, however the cultivated diversity using the approaches applied here reared a collection of limited diversity. It is likely that additional media types and isolation strategies could result in additional cultivated diversity as there are a number of taxa with aerobic heterotrophic lifestyles in the Core80 that have been brought into pure culture (e.g., *Microbulbifer*, *Hoeflea*). One challenge we experienced using the nutrient replete media was overgrowth of plates, even at 10 °C.

### 3.2 Secondary metabolite distributions and bioaccumulation in marine biota

Although the results of the archipelago spatial ascidian survey did not support a direct relationship between PalA levels and the relative abundance of microbiome ASVs, the results of the PalA niche analysis suggests that the Core80 ASVs occur in a preferred optimum and tolerance range of PalA levels. The lack of specific ASV-PalA patterns may not be entirely surprising, as secondary metabolites result from a complex combination of metabolic reactions that require a fine-tuning to environmental conditions and further metabolic modeling for the sake of understanding. Furthermore, these metabolites have been found to accumulate in the tissues in several different marine invertebrates. The Optimal Defense Theory can be applied to marine invertebrates and reflects the hypothesis that secondary metabolites are distributed in specific tissues based on exposure and anatomic susceptibility for predation [40]. For example, nudibranchs sequester toxic compounds, which have been biosynthesized by the gastropod or acquired from their prey. The toxins are concentrated in the anatomical space of their mantles, the most vulnerable portion of their soft, exposed bodies [40–42]. Bioaccumulation of secondary metabolites in invertebrates with less anatomical differentiation is also known to occur. In the phylum Porifera, different cell types and layers have been studied to determine spatial and anatomical differences in secondary metabolite concentrations [43–45]. Compounds have been found to be concentrated spatially on the surface (e.g., [46]) or apical parts of the sponge [47] in some cases. Sponges may be able to differentially bioaccumulate secondary cytotoxic metabolites based on tissues more susceptible to predation [48]. Metabolite distribution investigations that are ascidian-specific are less well documented; however, there is also evidence of ascidian secondary cytotoxic metabolite bioaccumulation. The patellazoles, marine macrolides from the ascidian *Lissoclinum patella*, bioaccumulate in the ascidian tissues to concentrations up to seven orders of magnitude higher than their cytotoxic dose in mammalian cell lines [49, 50]. Additionally, there are other instances in which bioaccumulation in ascidian host tissues suggests metabolic cooperation of producer and host as well as compound translocation from producer to host [15, 51, 52]. Although the PalA levels were normalized to grams of dry lobe weight, tissue-specific spatial localization is a potentially confounding factor in the statistical analyses investigating the ASV:PalA relationship.

### 3.3 Biosynthetic potential of the core

We investigated the natural product biosynthetic potential of the nine genera associated with 15 of 21 Core80 ASVs using antiSMASH (Table 2, Table S3). From this, it appears that all genera had at least one relative at the genus level with biosynthetic capacity for either polyketide or nonribosomal peptide biosynthesis or both. Even though the number of genomes available to survey were highly uneven, there is quite a disparity of biosynthetic capacity between the genera analyzed, thus, it appears that *Pseudovibrio*, *Nitrosomonas*, *Microbulbifer*, *Nitrospira* have the greatest capacities (in that order). Likewise, *Microbulbifer*, *Pseudovibrio*, *Hoeflea* and Opitutaceae might be prioritized as candidate PalA producers based solely on relative abundance ranking (Table 2; [24]). Although we did not conduct this analysis for the six ASVs that were classified at best at the family or order level, a few of these might be worth considering as potential producers considering their higher-level relationships with marine natural product producing lineages. For example, marine actinobacteria are classically associated with the production of numerous bioactive natural products (e.g., [53, 54]), although speculation is difficult with actinobacteria SaM_ASV20 in the core as it is only distantly related to known natural product producers. Likewise, Opitutaceae-related SaM_ASV15 is ranked 7 in terms of average relative abundance and falls in the same family the ascidian-associated *Candidatus* Didemnitutus mandela, which harbors the biosynthetic gene cluster predicted to produce mandelalide, a glycosylated polyketide [55]. From this we might prioritize the *Microbulbifer*, *Pseudovibrio*, and Opitutaceae ASVs for downstream investigation, with the lower relative abundance *Nitrosomonas* and *Nitrospira* ASVs also holding some potential given the perhaps surprising abundance of biosynthetic gene cluster content in these chemoautotrophic, and generally small genome size taxa. Although this analysis focused on predicted pathway characterizations across genera detected, the distributions of predicted pathways varied substantially across the taxa analyzed. The potential for these new Antarctic ascidian-associated strains to harbor secondary metabolite pathways remains speculative as they are amongst the most variable component of a bacterium's genome.

**Table 2.**
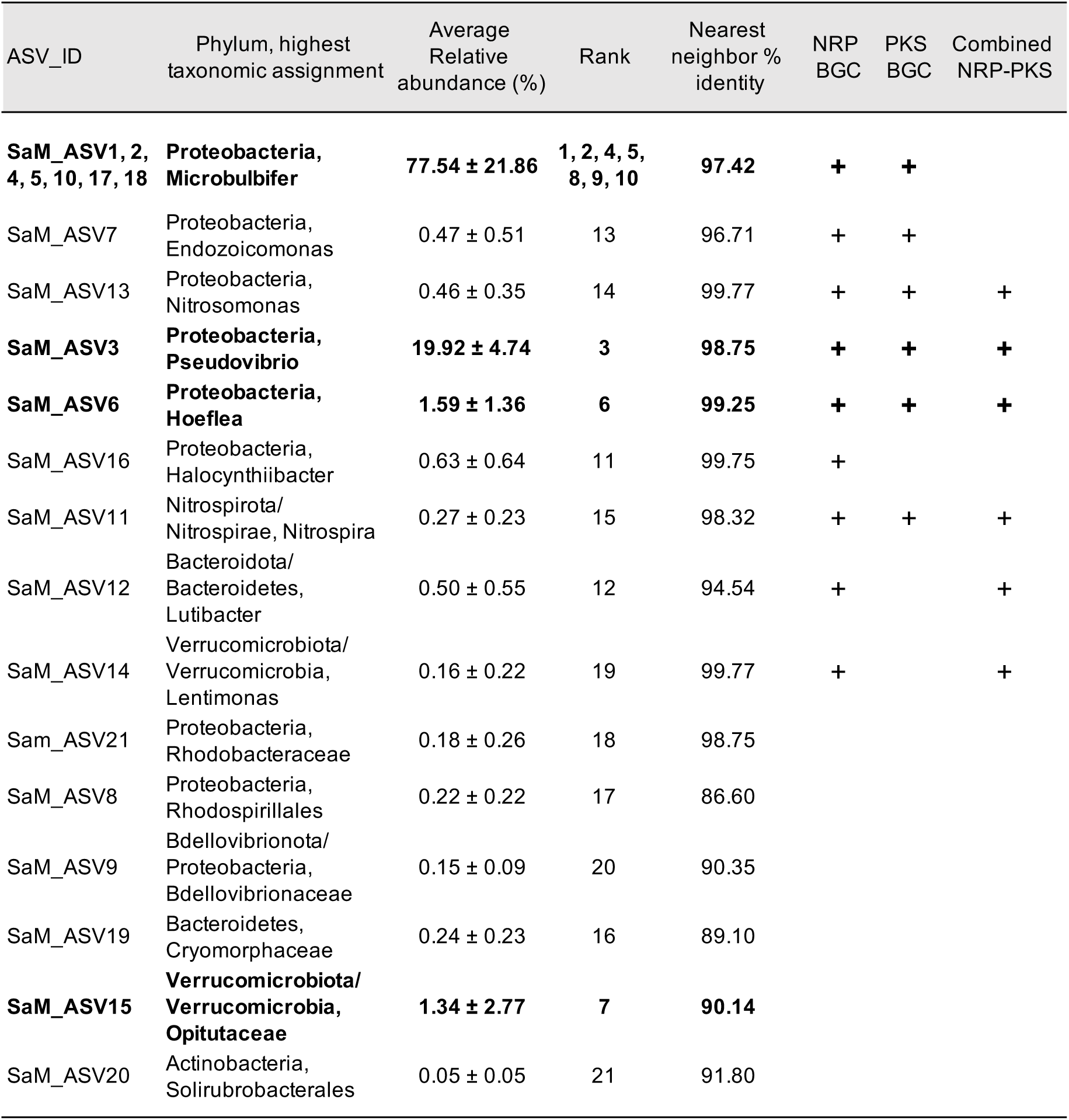
Taxonomic affiliations of core microbiome, relative abundance rank, and potential of affiliated genus in natural product gene cluster biosynthesis. Taxonomy is shown according to genome taxonomy database (GTDB) classification and NCBI taxonomy is included (GTDB/NCBI) where they differ. Biosynthetic potential only calculated for ASVs with Genus-level taxonomic assignments was based on representative genome biosynthetic gene cluster content in the same genus (See Figure S3 for list of genomes). ASVs in bold ranked in the top 10. Where more than one ASV was found per genus, the average relative abundance and standard deviations were summed. n=63 individuals.

## 4. Materials and Methods

### 4.1 Cultivation-dependent effort

*S. adareanum* samples collected by SCUBA in 2004 and 2007 were used for cultivation (**Table S4**). The 2004 specimen were archived in 20% glycerol at −80 °C until processing by manual homogenization using sterilized mortar and pestle prior to plating a suspension onto marine agar 2216 plates in 2006. The 2007 samples were homogenized immediately following collection using sterilized mortar and pestle, and suspensions were prepared in 1X marine broth or filter-sterilized seawater then transferred at 4 °C to DRI. Shortly after (within 1 month of collection), the 2007 isolates were cultivated on three types of media in which suspensions initially stored in marine broth were plated onto marine agar (2216), while homogenate preparations stored in seawater were plated onto VNSS agar media [56] and amended seawater plates (3 grams yeast extract (Difco), 5 g peptone (Difco), and 0.2 grams casein hydroslyate (Difco) per liter). Colonies were selected from initial plates, and purified through three rounds of growth on the same media they were isolated on.

### 4.2 Field sample collections for cultivation-independent efforts

Next, a spatial survey of *Synoicum adareanum* was executed in which samples were collected by SCUBA in austral fall between 23 March and 3 April 2011. Seven sampling sites (depths 24.7 - 31 m; Table S5) around the region accessible by Zodiac boat from Palmer Station were selected in which we sampled in a nested design where three multi-lobed colonies were selected from each site, and three lobes per colony were sampled (Figure 1). Underwater video [57] was taken at each site, then video footage was observed to note general ecosystem characteristics (% cover of major benthic species and algae). In total, 63 *S. adareanum* lobes were sampled (9 from each site). Samples were transported to Palmer Station on ice, and frozen at −80 °C until processing at DRI and USF. Frozen *S. adareanum* lobes were cut longitudinally in half for parallel processing through DNA and palmerolide detection pipelines.

Then, to address whether the composition of the SaM was distinct from the free-living bacterioplankton (< 2.5-μm fraction), we considered the overlap in membership between the SaM and bacterioplankton in the water column. To accomplish this, we used a reference seawater data set represented by samples that had been collected in February-March 2008 (five samples) and in August-September 2008 (nine samples) from LTER Station B near Anvers Island (east of the Bonaparte Point dive site), and at a few other locations in the region (Table S6). Seawater samples were collected by a submersible pump and acid washed silicone tubing at 10 m at Station B, and using a rosette equipped with 12 L Niskin bottles for the offshore samples at 10 m and 500 m (2 samples each depth). The February-March seawater samples were processed using in-line filtration with a 2.5 μm filter (Polygard, Millipore) to screen larger organisms and bacterioplankton were concentrated using a tangential flow filtration system and the cells were harvested on 25 mm 0.2 μm Supor filters (Millipore). The August-September seawater samples were processed using inline filtration with a 3.0 μm filter (Versapor, Millipore), and then bacterioplankton was collected onto 0.2 μm Sterivex (Millipore) filters using a multichannel peristaltic pump (Masterflex). All filters were immersed in sucrose:Tris:EDTA buffer [58] and stored frozen at −80 °C until extraction.

### 4.3 Palmerolide A screening

Sixty-three frozen *Synoicum adareanum* lobes were cut in half. Half lobes were lyophilized and then exhaustively extracted using dichloromethane for three days, followed by methanol for three days. The extracts were combined and dried on a rotary evaporator. The extracts were filtered, dried down, and reconstituted at 1.0 mg/mL to ensure the injected concentration was consistent. The residue was subjected to Liquid Chromatography-Mass Spectrometry (LC-MS) analysis using a H_2_O:ACN gradient with constant 0.05% formic acid. The high-resolution mass spectra were recorded on an Agilent Technologies 6230 electrospray ionization Time-of-Flight (ESI-ToF) spectrometer. LC-MS was performed using a C-18 Kinetex analytical column (50 x 2.1 mm; Phenomenex). The presence of PalA was verified using MS/MS on an Agilent Technologies 6540 UHD Accurate-Mass QTOF LC-MS.

The *S. adareanum*-associated microbial culture collection was screened for the presence of PalA. Isolates were cultivated in 30 mL volumes, and the resulting biomass was lyophilized and screened by HPLC. Each sample was analyzed in ESI-SIM mode targeting masses of 585 amu and 607 amu. The reversed-phase chromatographic analysis consisted of a 0.7 mL/min solvent gradient from 80% H_2_O:CH_3_CN to 100% CH_3_CN with constant 0.05% formic acid using the same C-18 column as above. The analysis was conducted for over 18 min. PalA had a retention time of approximately 14 min. The twenty-four sample sequence was followed by a PalA standard to confirm its analytical characteristics.

### 4.4 S. adareanum-associated microbial cell preparation

The outer 1-2 mm of the ascidian tissue, which could contain surface-associated microorganisms, was removed using a sterilized scalpel before sectioning tissue subsamples (~0.2 g) of frozen (−80 °C) *S. adareanum* (½ lobe sections). Tissue samples were diced with a scalpel before homogenization in sterile autoclaved and filtered seawater (1 mL) in 2 mL tubes. Each sample was homogenized (MiniLys, Bertin Instruments) using sterile CK28 beads (Precellys, Bertin Instruments) 3X at 5000 rpm for 20 sec, samples were placed on ice in between each homogenization. Homogenates were centrifuged at 500 × g at 4 °C for 5 min to pellet tissue debris. The supernatant was removed to a new tube for a second spin at the same conditions. This supernatant was decanted and the cell suspension centrifuged at 12,000 × g at 4 °C for 5 min to collect the microbial cells. Suspensions were stored on ice, then entered an extraction pipeline in which 12 samples were processed in parallel on the QIAvac 24 Plus Manifold (Qiagen).

### 4.5 DNA extractions

*S. adareanum*-associated microbial cell preparations were extracted with Powerlyzer DNEasy extraction (Qiagen) following manufacturer's instructions starting at with the addition of the lysis solution. Samples were processed in parallel in batches of twelve at a time using the QiaVac 24 Plus Vacuum Manifold (Qiagen). The lysis step (2X at 5000 rpm for 60 seconds each with incubation on ice was performed on the MiniLys (Precellys, Bertin Instruments) using the 0.1 mm glass beads that come with the Powerlyzer kit. DNA concentrations of final preparations were estimated using Quant-iT Picogreen dsDNA Assay Kit (Invitrogen) fluorescence detection on a Spectramax Gemini (Molecular Devices).

DNA from bacterioplankton samples was extracted following [58], and DNA from bacterial cultures was extracted using the DNeasy Blood and Tissue kit (Qiagen) following manufacturer's instructions. All DNA concentrations were estimated using Picogreen.

### 4.6 16S rRNA gene sequencing

Illumina tag sequencing for the *S. adareanum* microbiome (SaM) targeted the V3-V4 region of the 16S rRNA gene using primers 341F CCTACGGGNBGCASCAG and 806R GGACTACHVGGGTWTCTAAT. The first round of PCR amplified the V3-V4 region using HIFI HotStart Ready Mix (Kapa Biosystems). The first round of PCR used a denaturation temperature of 95 °C for 3 min, 20 cycles of 95 °C for 30 sec, 55 °C for 30 sec and 72 °C for 30 sec and followed by an extension of 72 °C for 5 minutes before holding at 4 °C. The second round of PCR added Illumina-specific sequencing adapter sequences and unique indexes, permitting multiplexing, using the Nextera XT Index Kit v2 (Illumina) and HIFI HotStart Ready Mix (Kapa Biosystems). The second round of PCR used a denaturation temperature of 95 °C for 3 minutes, 8 cycles of 95 ºC for 30 seconds, 55 °C for 30 seconds and 72 °C for 30 seconds and followed by an extension of 72 °C for 5 minutes before holding at 4 °C. Amplicons were cleaned up using AMPure XP beads (Beckman Coulter). A no-template control was processed but did not show a band in the V3-V4 amplicon region and was sequenced for confirmation. A Qubit dsDNA HS Assay (ThermoFisher Scientific) was used for DNA concentration estimates. The average size of the library was determined by the High Sensitivity DNA Kit (Agilent) and the Library Quantification Kit – Illumina/Universal Kit (KAPA Biosystems) quantified the prepared libraries. The amplicon pool sequenced on Illumina MiSeq generated paired end 301 bp reads was demultiplexed using Illumina's bcl2fastq.

The bacterioplankton samples sent to the Joint Genome Institute (JGI) for library preparation and paired end (2 x 250 bp) MiSeq Illumina sequencing of the variable region 4 (V4) using primers 515F (5′-GTGCCAGCMGCCGCGGTAA-3′) and 806R (5′-GGACTACHVGGGTWTCTAAT-3′) [59].

Sequence processing included removal of PhiX contaminants and Illumina adapters at JGI. The identity of the cultivated isolates was confirmed by16S rRNA gene sequencing using

Bact27F and Bact1492R primers either by directly sequencing agarose-gel purified PCR products (Qiagen), or TA cloning (Invitrogen) of PCR fragments into *E. coli* following manufacturer's instructions in which three clones were sequenced for each library, plasmids were purified (Qiagen) at the Nevada Genomics Center, where Sanger sequencing was conducted on an ABI3700 (Applied Biosystems). Sequences were trimmed and quality checked using Sequencer, v. 5.1.

### 4.7 16S rRNA gene sequencing

We employed a QIIME2 pipeline [60] using the DADA2 plug-in [61] to de-noise the data and generate amplicon sequence variant (ASV) occurrence matrices for the SaM and bacterioplankton samples. The rigor of ASV determination was used in this instance given the increased ability to uncover variability in the limited geographic study area, interest in uncovering patterns of host-specificity, and ultimately in identifying the conserved, core members of the microbiome, at least one of which may be capable of PalA biosynthesis. Sequence data sets were initially imported into QIIME2 working format and the quality of forward and reverse were checked. Default trimming parameters included trimming all bases after the first quality score of 2, in addition, the first 10 bases were trimmed, and reads shorter than 250 bases were discarded. Next the DADA2 algorithm was used to de-noise the reads (corrects substitution and insertion/deletion errors and infers sequence variants). After de-noising, reads were merged. The ASVs were constructed by grouping the unique full de-noised sequences (the equivalent of 100% OTUs, operational taxonomic units). The ASVs were further curated in the QIIME2-DADA2 pipeline by removing chimeras in each sample individually if they can be exactly reconstructed by combining a left-segment and a right-segment from two more abundant “parent” sequences. A pre-trained SILVA 132 99% 16S rRNA Naive Bayes classifier (https://data.qiime2.org/2019.1/common/silva-132-99-nb-classifier.qza) was used to perform the taxonomic classification. Compositions of the SaM and the bacterioplankton ASVs were summarized by proportion at different taxonomy levels, including genus, family, order, class, and phylum ranks. In order to retain all samples for diversity analysis, we set lowest reads frequency per sample (n = 62 samples at 19003 reads; n = 63 samples at 9987 reads) as rarefaction depth to normalize the data for differences in sequence count. ASVs assigned to Eukarya or with unassigned taxa (suspected contaminants) were removed from the final occurrence matrix such that the final matrix read counts were slightly uneven with the lowest number of reads per sample with 9961 reads.

The SaM ASVs were binned into Core (highly persistent) if present in ≥80% of samples (Core80), Dynamic if present in 50-79% of samples (Dynamic50) and those that comprise the naturally fluctuating microbiome, or Variable fraction that was defined as those ASVs present in <50% of the samples [2, 3]. We used these conservative groupings of the core microbiome due to the low depth of sequencing in our study [3].

ASV identities between the SaM, the *S. adareanum* bacterial isolates and the bacterioplankton data sets were compared using CD-HIT (cd-hit-est-2d; http://cd-hit.org). The larger SaM data set which included 19,003 sequences per sample was used for these comparisons to maximize the ability to identify matches; note that this set does exclude one sample, Bon1b which had half as many ASVs, though overall this larger data set includes nearly 200 additional sequences in the Variable fraction for comparison. ASVs with 100 and 97% identity between the pairwise comparisons were summarized in terms of their membership in the Core, Dynamic or Variable fractions of the SaM. Likewise, CD-HIT was used to dereplicate the isolate sequences at a level of 99% sequence identity, and then the dereplicated set was compared against the bacterioplankton iTag data set.

Phylogenetic analysis of the SaM ASVs, *S. adareanum* bacterial isolates, and 16S rRNA gene cloned sequences from Riesenfeld et al. [13] was conducted with respect to neighboring sequences identified in the Ribosomal Database Project and SILVA and an archaea outgroup using MEGA v.7 [62]. Two maximum likelihood trees were constructed; the first with the Core80 ASVs, and the second with both Core80 and Dynamic50 ASVs. A total of 369 aligned positions were used in both trees. A total of 1000 bootstrap replicates were run in both instances, in which the percentage (≥ 50 %) of trees in which the associated taxa clustered together are shown next to the branches.

### 4.8 Statistical analyses

T-tests were run in Statistica (v. 13) to determine significance (p < 0.05) of site-to-site, within and between colony variation in PalA concentrations. Similarity-matrix and hierarchical clustering analyses were performed using PRIMER v.7 and PERMANOVA+ (PRIMER-e). Analyses were performed on the complete microbiome as well as the three microbiome fractions in most cases. The ASV occurrence data was square root transformed for all analyses. A heat map based on the Core80 ASV occurrence was generated with the transformed data, and hierarchical clustering with group average parameter was employed, which was integrated with SIMPROF confidence using 9,999 permutations and a 5% significance level. Bray-Curtis resemblance matrixes were created using ASV occurrences without the use of a dummy variable. To determine basic patterns in community structure within vs. between colony and site differences the significance was determined by t-test using Statistica v. 13. To compare within colony (n = 9) vs. between colony differences (n = 27), nine “between colony pairwise similarity values were randomly sampled in order to compare equal sample sizes, checking that the homogeneity of variance was similar between them. Then threshold metric Multi-Dimensional Scaling (tmMDS) was conducted based on Kruskal fit scheme 1, including 500 iterations, and a minimum stress of 0.001. Similarity profile testing through SIMPROF were performed based on a null hypothesis that no groups would demonstrate differences in ASV occurrences. This clustering algorithm was also used to generate confidence levels on the MDS plot, which were set to 65% and 75%. In addition, 95% bootstrap regions were calculated with 43 bootstraps per group, set to ensure a minimum rho of 0.99. In order to assess the contribution of each factor to the variance of the microbial community in this nested experimental design, Site(Colony), permutational multivariate analysis of variance (PERMANOVA) was used. Site-based centroids were calculated and the PERMDISP algorithm was used to determine the degree of dispersion around the centroid for each site. Overall site-to-site difference in dispersion was determined and pairwise comparisons were also calculated with 9,999 permutations used to determine significance (P(perm) <0.05). Exploratory analysis of the major ASV contributors to similarity was performed using the SIMPER procedure based on sites and colonies, with a cut off for low contributions set to 70%.

Co-occurrence networks were constructed using filtered ASV occurrence data sets in which the ASV were filtered to only those that were present in at least five samples resulting in a 102 ASV data set. The 102 x 63 matrix was provided as input to FlashWeave v1.0 [63] using default parameters, and visualized in Gephi v. 0.9.2 [64]. Then to consider whether the ASVs in the Core80, Dynamic50, or Variable fractions of the SaM were affiliated with particular levels of PalA in the ascidian lobes, PalA niche robust optimum and range were computed using the occurrence and dry weight normalized contextual data [65]. Weighted gene correlation network analysis (WGCNA package in R [66]) was used to identify modules and their correlation with PalA levels. The matrix was total-sum normalized [67], and WGCNA was used in signed mode. There were few modules detected, although they were not correlated with PalA. Modules were projected on the FlashWeave co-occurrence network and called subsystems.

### 4.9 Biosynthetic gene cluster analysis

Subsequently, in order to predict the likelihood of Core80 ASV lineages harboring the potential for natural product biosynthesis we designed a meta-analysis of neighboring genomes found at the Integrated Microbial Genomes (IMG) database [68]. The analysis was conducted only for ASVs in which confidence of taxonomic assignment was at the genus level. Therefore, genomes were harvested from IMG that were associated with a total of 9 genera (*Microbulbifer* (16 genomes), *Pseudovibrio* (24 genomes), *Endozoicomonas* (11 genomes), *Nitrosomonas* (19 of 68 total genomes in this genus), *Nitrospira* (14 genomes), *Hoeflea* (7 genomes), *Lutibacter* (12 genomes), *Halocynthilibacter* (2 genomes), and in the case of *Lentimonas*, since no genomes were found, we harvested 8 genomes from the Puniciococcaceae family). This results in 113 genomes that were submitted to antiSMASH [69] for analysis. The genomes and counts of biosynthetic gene clusters assigned to nonribosomal peptide synthase, polyketide synthase, or hybrid of the two classes were tabulated (Table S2).

### 4.10 Data availability

*Synoicum adareanum* microbiome Illumina sequence information and associated metadata are described under NCBI BioProject PRJNA597083 (https://www.ncbi.nlm.nih.gov/bioproject/?term=PRJNA597083), and the *S. adareanum* culture collection 16S rRNA gene sequences were deposited in GenBank under MN960541-MN960556. The bacterioplankton sequence information and associated metadata are described under NCBI BioProject PRJNA602715 (https://www.ncbi.nlm.nih.gov/bioproject/?term=PRJNA602715). Records for these Antarctic metadata and associated sequences depositions are also reflected in the mARS database (http://mars.biodiversity.aq).

## 5. Conclusions

This work has advanced our understanding of the Antarctic ascidian *S. adareanum*, PalA distributions, and microbiome in several ways. First, we found PalA to be a dominant product across all 63 samples, with some variation but no coherent trends with the site, sample, or microbiome ASV. The results point to a conserved, core, microbiome represented by 21 ASVs, 20 of which appear to be distinct from the bacterioplankton. The phylogenetic distribution of these taxa was diverse, and distinct from other ascidian microbiomes in which organisms with both heterotrophic and chemosynthetic lifestyles are predicted. The co-occurrence analysis suggested the potential for ecologically interacting microbial networks that may improve our understanding of this ascidian-microbiome-natural product system. Likewise, based on the occurrence of natural product biosynthetic gene clusters, there are several potential PalA producers. These results advance the long-term goal of *Synoicum adareanum*-palmerolide-microbiome research which is compelled by the fact that by identifying the producer, genome sequencing could then provide information on PalA biosynthesis which could lead to the development of a potential therapeutic agent to fight melanoma.

## Supplementary Materials

The following are available online at www.mdpi.com/xxx/s1, Table S1: PERMANOVA estimators of drivers of variability, Table S2: Taxonomic distribution of bacterioplankton ASVs, Table S3 Biosynthetic gene clusters in bacterial genomes related to Core80 SaM genera, Table S4: *Synoicum adareanum* collections and preparations for microbiome cultivation, Table S5 *Synoicum adareanum* collections for palmerolide A and microbiome characterization by V3-4 rRNA gene tag sequencing, Table S6: Bacterioplankton collections used in v4 rRNA gene tag sequencing. Figure S1: Maximum likelihood 16S rRNA gene phylogenetic tree, Figure S2: Results of pairwise t-tests of PalA levels determined by mass spectrometry, Figure S3: Average pairwise similarity within and between *S. adareanum* microbiome community structures, Figure S4: tmMDS plots representing the microbiome of the 63 *S. adareanum* samples, Figure S5. PalA niche optimum for *S. adareanum* microbiome ASVs.

## Author Contributions

This work was the result of a team effort in which the following contributions are recognized: conceptualization, A.E.M., P.S.G.C, and B.J.B. methodology and experimentation, A.E.M., N.E.A., L.B., D. E., M.L.H., S.K., C.S.R. and R.M.Y.; validation, K.D., M.L.H., and B.J.B. formal analysis, A.E.M., N.E.A., E.D., D.E., S.K., and C-C.L.; data curation, M.L.H. and C-C.L.; writing—original draft preparation, A.E.M., B.J.B., N.E.A., D.E., S.K., and C-C.L.; writing—review and editing, A.E.M., N.E.A, B.J.B., L.B., P.S.G.C., A.E.K.D., D.E., S.K., C.S.R., and R.M.Y.; supervision, A.E.M., B.J.B., P.S.G.C., K.W.D., A.E.K.D., and C.S.R.; project administration, K.W.D.; funding acquisition, A.E.M., P.S.G.C., and B.J.B. All authors have read and agreed to the published version of the manuscript.

## Acknowledgments

The assistance of several collaborators and students is also acknowledged including Charles Amsler, Margaret Amsler, Jason Cuce, Bill Dent, Alex Dussaq, Cheryl Gleasner, Alan Maschek, Robert W. Read, Andrew Shilling, Santana Thomas, and the Palmer Station science support staff.

