## Supplementary tables and figures for "Uncovering the core microbiome and distributions of palmerolide in *Synoicum adareanum* across the Anvers Island archipelago, Antarctica"

**Supplementary Information**

Table S1. PERMANOVA estimators of drivers of variability. This allows the components of the nested experimental design, site and colony, to be evaluated. Note, all values presented in the table are statistically significant (P(perm)< 0.05).

|  | All ASVs (%) | Core (%) | Dynamic (%) | Variable (%) |
| --- | --- | --- | --- | --- |
| Site-to-Site | 25.0 | 26.2 | 28.4 | 19.2 |
| Colony-to-Colony | 27.8 | 31.5 | 27.2 | 22.3 |
| Residual | 47.2 | 42.3 | 44.4 | 58.4 |

Table S2. Taxonomic distribution of bacterioplankton ASVs. Phyla and assigned classes are indicated, along with average representation across the 15-sample data set. Average values below 0.005 are represented as 0.00%.

| Phylum_Class | Average Representation |
| --- | --- |
| Acidobacteria_Acidobacteriia | 0.00% |
| Acidobacteria_Subgroup 26 | 0.00% |
| Acidobacteria_Subgroup 6 | 0.01% |
| Actinobacteria_Acidimicrobiia | 0.96% |
| Actinobacteria_Actinobacteria | 0.05% |
| AncK6_Unclassified | 0.01% |
| Atribacteria_JS1 | 0.00% |
| Bacteroidetes_Bacteroidia | 13.10% |
| Chlamydiae | 0.03% |
| Chloroflexi_Dehalococcoidia | 0.49% |
| Chloroflexi_JG30-KF-CM66 | 0.03% |
| Oxyphotobacteria/Plastid | 7.39% |
| Dadabacteria_Dadabacteriia | 0.06% |
| Dependentiae_Babeliae | 0.00% |
| Epsilonbacteraeota_Campylobacteria | 0.32% |
| Fibrobacteres_Chitinivibrionia | 0.00% |
| Fibrobacteres_Fibrobacteria | 0.00% |
| Firmicutes_Bacilli | 0.03% |
| Firmicutes_Clostridia | 0.01% |
| Fusobacteria_Fusobacteriia | 0.03% |
| Gemmatimonadetes_BD2-11 terrestrial group | 0.12% |
| Hydrogenedentes_Hydrogenedentia | 0.00% |
| Kiritimatiellaeota_Kiritimatiellae | 0.00% |
| Latescibacteria_Latescibacteria | 0.00% |
| Lentisphaerae_Lentisphaeria | 0.01% |
| Lentisphaerae_Oligosphaeria | 0.00% |
| Lentisphaerae_Unclassified | 0.00% |
| Margulisbacteria_Unclassified | 0.04% |
| Marinimicrobia (SAR406 clade)_Unclassified | 2.72% |
| Nitrospinae_Nitrospinia | 0.31% |
| Nitrospinae_P9X2b3D02 | 0.00% |
| Nitrospinae_Nitrospira | 0.00% |
| Unclassified_Omnitrophicaeota | 0.00% |
| Patescibacteria_Gracilibacteria | 0.03% |
| Patescibacteria_Parcubacteria | 0.00% |
| PAUC34f_Unclassified | 0.13% |
| Planctomycetes_BD7-11 | 0.01% |
| Planctomycetes_Brocadiae | 0.00% |
| Planctomycetes_OM190 | 0.03% |
| Planctomycetes_Phycisphaerae | 0.10% |
| Planctomycetes_Pla3 lineage | 0.01% |
| Planctomycetes_Pla4 lineage | 0.00% |
| Planctomycetes_Planctomycetacia | 0.15% |
| Planctomycetes_SGST604 | 0.00% |
| Planktomycetes_unclassified | 0.00% |
| Poribacteria_Unclassified | 0.00% |
| Proteobacteria_Alphaproteobacteria | 15.57% |
| Proteobacteria_Deltaproteobacteria | 2.81% |
| Proteobacteria_Gammaproteobacteria | 47.35% |
| Proteobacteria_Unclassified | 0.00% |
| Stramenopiles | 0.00% |
| Spirochaetes_Spirochaetia | 0.00% |
| Tenericutes_Mollicutes | 0.00% |
| Verrucomicrobia_Verrucomicrobiae | 0.28% |
| Unclassified_Bacteria | 0.07% |
| Thaumarchaeota_Nitrososphaeria | 6.29% |
| Euryarchaeota_Halobacteria | 0.00% |
| Euryarchaeota_Thermoplasmata | 1.39% |
| Nanoarchaeota_Woesearchaeia | 0.00% |

Table S3. Biosynthetic gene clusters in bacterial genomes related to Core80 SaM genera. The biosynthetic gene clusters were identified using antiSMASH. Genome sequence identifiers are indicated (initially identified in the Integrated Microbial Genomes database, IMG) in addition to NCBI accession numbers, where possible, along with the number of gene clusters identified that were related to polyketide biosynthesis (PKS) or non-ribosomal peptide synthesis (NRPS).

Table S4. *Synoicum adareanum* collections and preparations for microbiome cultivation. Temperature was estimated using dive computer.

| *S. adareanum* sample ID | Collection Site | GPS Location | Date | Depth (m) | Temperature ( °C) |
| --- | --- | --- | --- | --- | --- |
| PSC04 | Norsel Point | S 64° 45.638'  W 64° 05.874' | 2004 | 28.3 | n.d. |
| P7-AM17 | Bonaparte Point | S 64° 46.662'  W 64° 03.986' | 2-May-07 | 23.4 | -1.1 |

Table S6. Bacterioplankton collections used in v4 rRNA gene tag sequencing.

| Picoplankton sample ID | Collection Site | GPS Location | Date | Depth (m) | Temperature ( °C) |
| --- | --- | --- | --- | --- | --- |
| Palmer PEL 1 2-22-08 | B-C | S 64° 47.009' W 64° 46.656' | 22-Feb-08 | 10 | 0.5 |
| Palmer PEL 2 2-25-08 | B-C | S 64° 47.009' W 64° 46.656' | 25-Feb-08 | 10 | 1.5 |
| Palmer PEL 4 2-2-08 | B-C | S 64° 47.009' W 64° 46.656' | 29-Feb-08 | 10 | 1.6 |
| Palmer PEL 5 3-3-08 | B-C | S 64° 47.009' W 64° 46.656' | 3-Mar-08 | 10 | 1.6 |
| Palmer PEL 6 3-6-08 | B-C | S 64° 47.009' W 64° 46.656' | 6-Mar-08 | 10 | 1.6 |
| IPY-206-3 | B | S 64° 46.769' W 64° 04.353' | 24-Jul-08 | 10 | -0.9 |
| IPY-210-4 | B | S 64° 46.769' W 64° 04.353' | 28-Jul-08 | 10 | -0.7 |
| IPY-225-9 | B | S 64° 46.769' W 64° 04.353' | 12-Aug-08 | 10 | -1.5 |
| IPY-243-14 | Palmer Deep | S 64° 55.556' W 64° 82.962' | 30-Aug-08 | 500 | -1.7 |
| IPY-243-15 | Palmer Deep | S 64° 55.556' W 64° 82.962' | 30-Aug-08 | 10 | -1.7 |
| IPY-245-16 | Gerlache Strait | S 64° 38.628' W 62° 53.343' | 1-Sep-08 | 500 | -0.7 |
| IPY-245-17 | Gerlache Strait | S 64° 38.628' W 62° 53.343' | 1-Sep-08 | 10 | -1.8 |
| IPY-246-19 | LaPeyrére Bay | S 64° 25.275' W 63° 16.974' | 2-Sep-08 | 10 | -1.1 |
| IPY-256-21 | B | S 64° 46.769' W 64° 04.353' | 12-Sep-08 | 10 | n.d. |

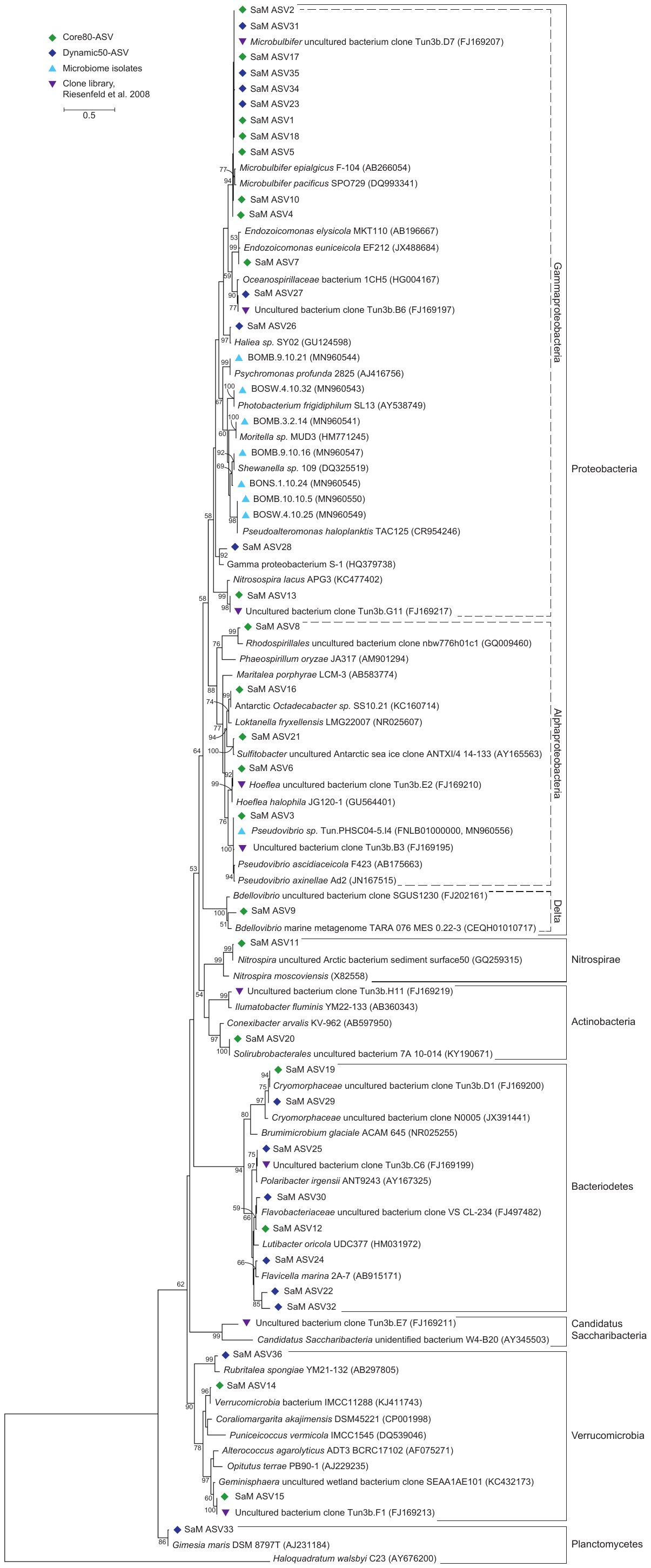

Figure S1. Maximum likelihood 16S rRNA gene phylogenetic tree (based on 369 aligned bases). Archaeon, *Haloquadratum walsbyi* was used as an outgroup. Core (green diamonds) and dynamic (blue diamonds) ASVs are shown, along with cultivated isolates from the *Synoicum adareanum* microbiome (light blue triangles) and cloned rRNA gene sequences from an earlier study with *S. adareanum* collected in 2006 [13]. *[note upon final submission, this figure will appear as a standalone PDF in which it will be in full size]*

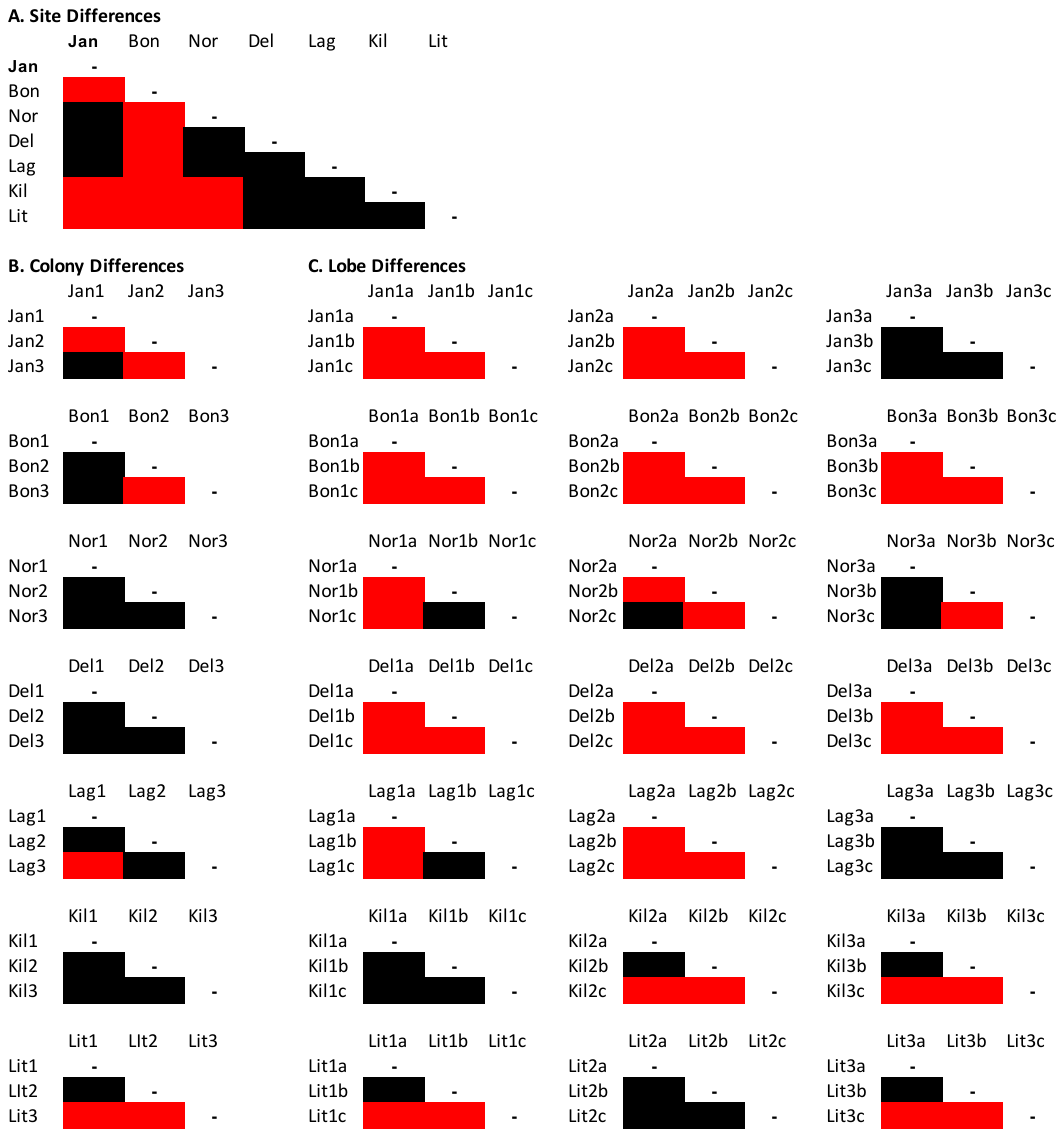

Figure S2. Results of pairwise t-tests of PalA levels determined by mass spectrometry in which significant (p ≤ 0.05) paired comparisons are indicated in red, and nonsignificant in black. n=9 for the site to site comparisons in (a); n=3 for colony to colony comparisons in (b); and n=3 for lobe:lobe comparisons in (c).

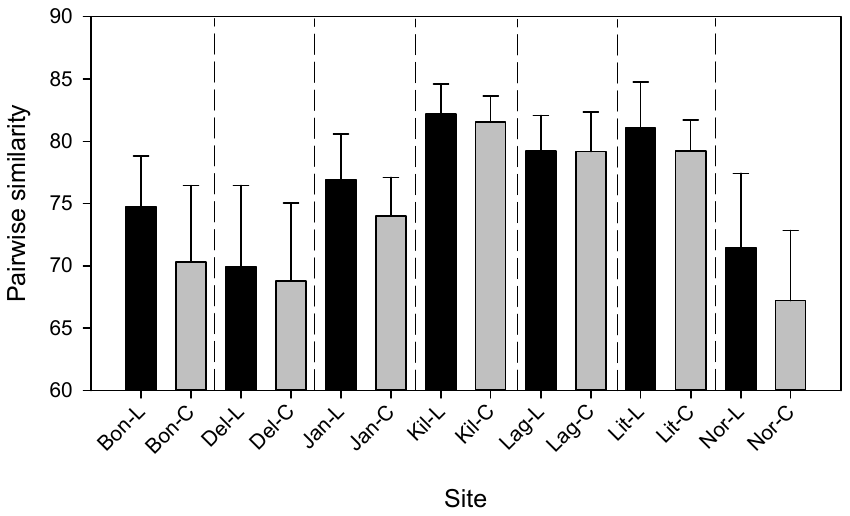

*

*

Figure S3. Average pairwise similarity within (labels with -L, black bars) and between *S. adareanum* colony microbiome community structures (labels with -C, gray bars) at each dive site (abbreviations for sites as in Figure 3). Standard deviation shown, n=9 for lobes and colonies (which were randomly sampled from 27). Bray Curtis similarity was calculated with square-root transformed occurrence data.

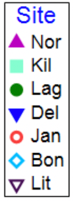

b

a

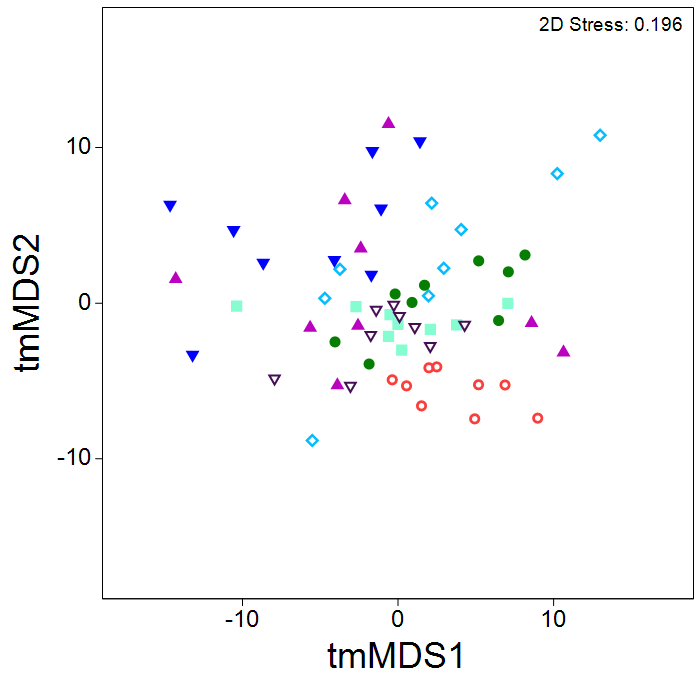

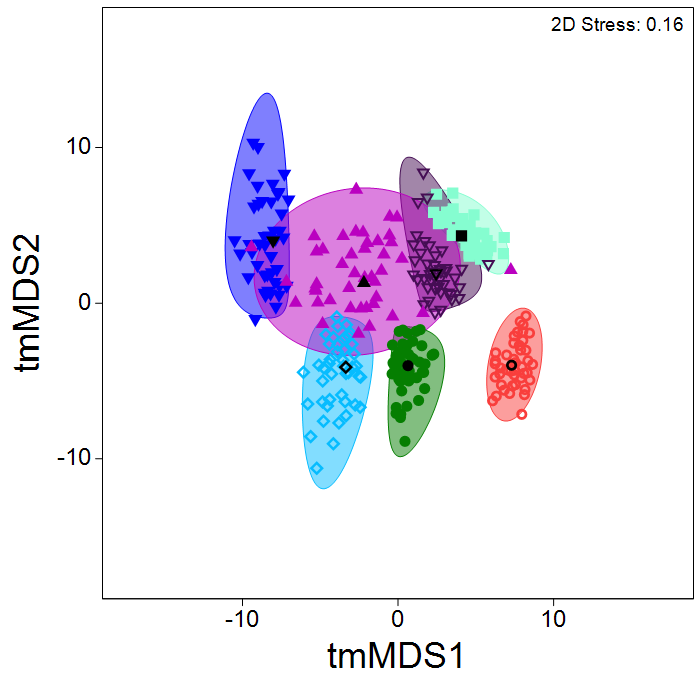

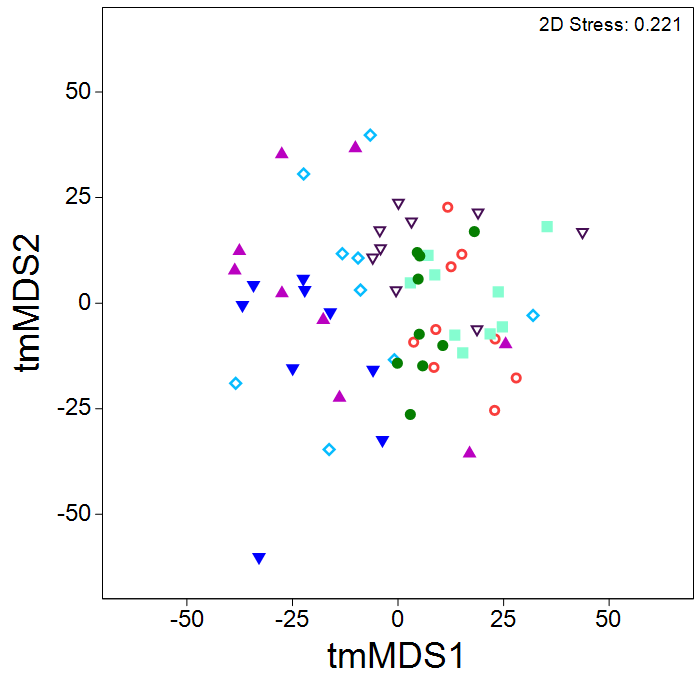

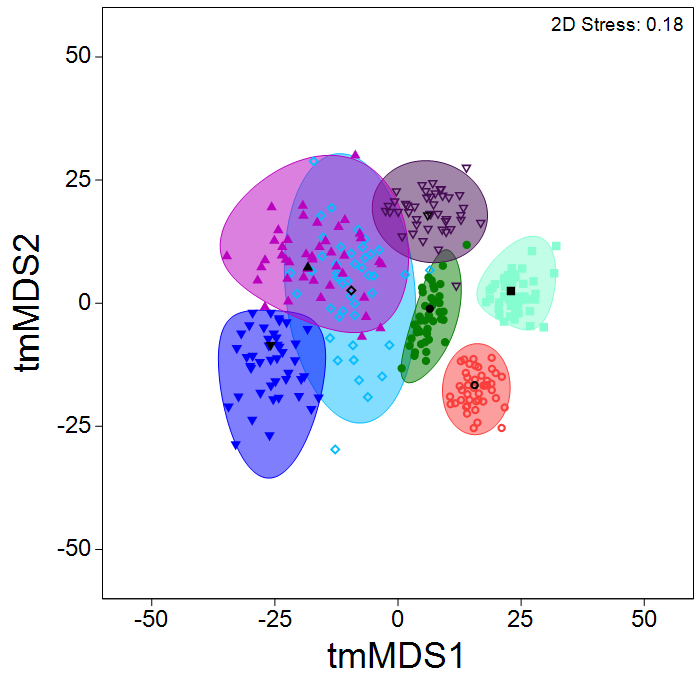

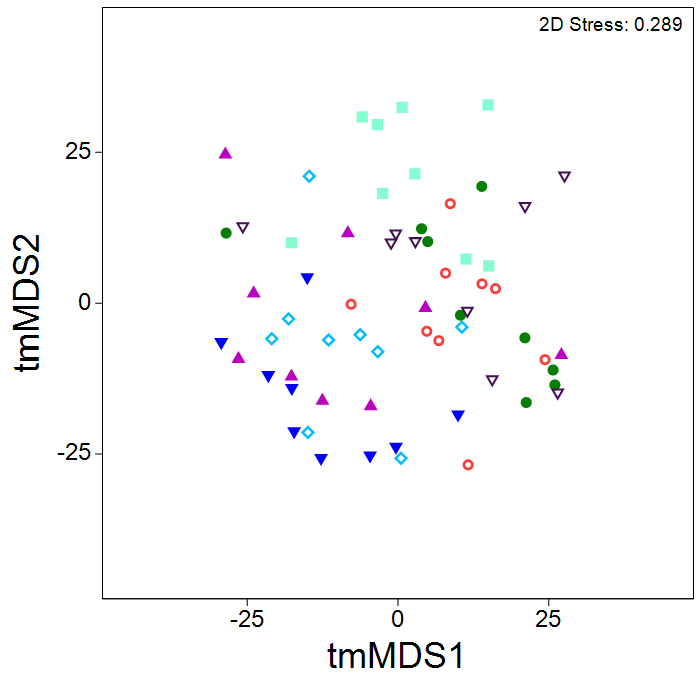

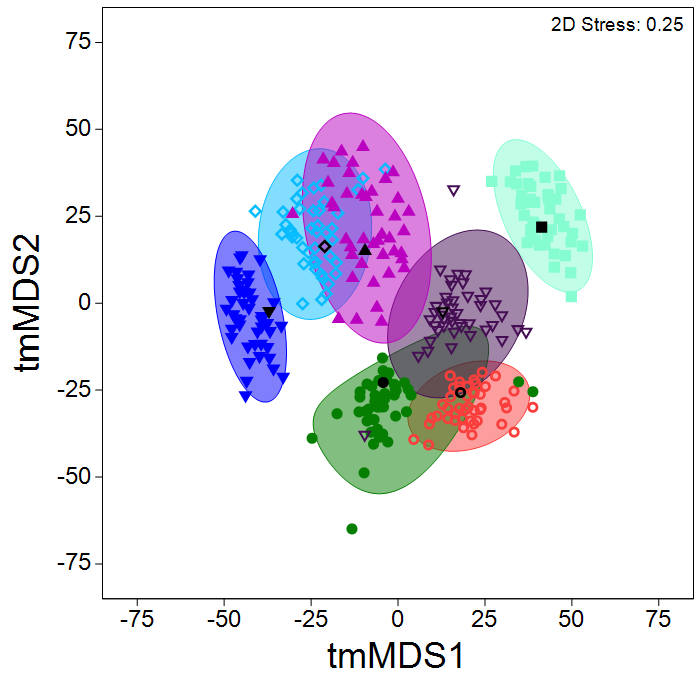

d

c

f

e

Figure S4. tmMDS plots (Bray-Curtis; square root transformed ASV occurrence data) representing the microbiome of the 63 *S. adareanum* samples using the ASV occurrence profiles for the core (a), dynamic (c), and variable (e) microbiome fractions. Bootstrapped data MDS plots (43 bootstraps) for subgroups are also provided (b, d, and f); the resulting data cloud for each site is shaded to aid visualization of the cluster. The site-based centroid is demarcated by its respective symbol in black.

Figure S5. PalA niche optimum for *S. adareanum* microbiome ASVs. PalA range and median values (diamond symbols) are shown according to microbiome membership (Core80, green; Dynamic50, blue; Variable, pink). Data were ordered from low to high PalA niche values (i.e. ASVs found in low PalA-containing tissues at the top, and those found associated with high PalA tissues at the bottom). Taxonomic phyla and highest level of assignment are indicated.
